# Monoamine-induced diacylglycerol signaling rapidly accumulates Unc13 in nanoclusters for fast presynaptic potentiation

**DOI:** 10.1101/2025.01.10.632340

**Authors:** Natalie Blaum, Tina Ghelani, Torsten Götz, Keagan S. Chronister, Mercedes Bengochea, Christian F. Christensen, Thiago C. Moulin, Livia Ceresnova, Hanna Kern, Ulrich Thomas, Martin Heine, Stephan J. Sigrist, Alexander M. Walter

## Abstract

Neuromodulators control mood, arousal, and behavior by inducing synaptic plasticity via G-protein coupled receptors. Long-term potentiation of presynaptic neurotransmitter release requires structural changes, but how fast potentiation is achieved within minutes remains enigmatic. Using the *Drosophila melanogaster* neuromuscular junction, we show that on the timescale of one minute, octopamine, the invertebrate analog of nor-epinephrine, rapidly potentiates evoked neurotransmitter release by a G protein coupled pathway involving presynaptic OAMB receptors and phospholipase C. No changes of presynaptic calcium influx were seen, but confocal signals of the release factor Unc13A and the scaffolding protein Bruchpilot increased within one minute of octopamine treatment. On the same timescale, live, single-molecule imaging of endogenously tagged Unc13 revealed its instantly reduced motility and its increased concentration in synaptic nanoclusters with potentiation. Presynaptic knockdown of Unc13A fully blocked fast potentiation and removal of its N-terminal localization sequence delocalized the protein fragment to the cytosol, but it was rapidly recruited to the plasma membrane by DAG analog phorbol esters and octopamine, pointing to a role in C-terminal domains. Point mutation of endogenous Unc13 disrupting diacylglycerol-binding to its C1 domain blocked plasticity-induced nanoscopic enrichment and synaptic potentiation. The mutation increased basal neurotransmission but reduced Unc13 levels, revealing a gain of function and potential homeostatic compensation. The mutation also blocked phorbol ester-induced potentiation, decreased the calcium-sensitivity of neurotransmission and caused short-term synaptic depression. At the organismal level, the mutation reduced locomotion and survival while enhancing reproduction. Thus, the Unc13 C1 domain mediates acute subsynaptic compaction of Unc13 under monoamine-induced potentiation and influences short-term plasticity, locomotion, reproduction, and survival.

## Introduction

Chemical synaptic transmission relies on the timed release of neurotransmitters from synaptic vesicles (SVs) in response to action-potential (AP) induced calcium influx through voltage gated ion channels to activate postsynaptic receptors. Synaptic plasticity, the change in the transmission strength, is needed across wide timescales for neural information processing, signal stabilization, and information storage. Postsynaptic plasticity adapts the responses neurotransmitters elicit on the receiving cell, while presynaptic plasticity alters their release.

Neurotransmitter release relies on a highly conserved molecular machinery to target (dock) SVs to the plasma membrane and to prime them molecularly such that they can fuse with the plasma membrane in response to an AP^1^. Docking, priming and SV fusion depend on (M)Unc13 and (M)Unc18 proteins (“M” in the mammalian case) and the assembly of the SNARE complex between the SV and plasma membrane proteins synaptobrevin, syntaxin-1 and SNAP25^2^. Neurotransmitter release is linked to the AP by the SV protein Synaptotagmin that triggers SV fusion in response to presynaptic calcium (Ca^2+^) influx through voltage gated ion channels^3^. The local topology of the synapse is controlled by large multi-domain cytomatrix proteins like the Rab3 Interacting Molecule (RIM), RIM-Binding Protein (RIM-BP) and Bruchpilot (BRP in *Drosophila melanogaster*) or ELKS/CAST (in mammals)^4, 5, 6^.

Long-term potentiation of neurotransmitter release is essential for homeostasis and information storage. It coincides with structural presynaptic adaptations, including local enrichment of Ca^2+^ channels, RIM, RIM-BP, BRP and (M)Unc13^7, 8, 9, 10, 11, 12, 13^. Among those factors, (M)Unc13 adaptations appear to be required, pointing to a pivotal role of the protein in this synaptic plasticity^7^. However, synapses must also adjust on shorter timescales (within 1 minute or less) to bridge the time it takes to manifest these changes. The mechanisms underlying intermittent potentiation remain unclear, but they may involve evolutionarily conserved (M)Unc13 domains CaM, C2B, and C1 which bind to Ca^2+^/Calmodulin, Ca^2+^/PI(4,5)P_2_ and DAG to regulate neurotransmitter release^14, 15^. Consistent with this, DAG analogs like phorbol esters rapidly potentiate neurotransmitter release via the C1 domain, though the reasons for this effect and the biological pathway involved remain elusive^16, 17^.

Neural G-protein coupled receptors (GPCRs) are major pharmacological targets in attempts to cure neurological disorders^18, 19^. Neuromodulator binding induces a conformational change leading to the intracellular dissociation of the heteromeric G-protein complex into the Gα and Gβ/γ subunits which modulate downstream effector pathways. These are distinguished into Gαi/o, Gαs, Gαq/11 and Gα12/13. Gαi/o and Gαs pathways inhibit or activate adenylate cyclase to generate the second messenger cyclic-adenosine monophosphate (cAMP) which activates Protein Kinase A (PKA) and regulates synapse growth, short-term plasticity, long-term potentiation, animal behavior, and learning and memory^20, 21, 22, 23, 24, 25, 26, 27, 28, 29^. The Gαq/11 pathway activates phospholipase C (PLC) to hydrolyze phosphatidylinositol 4,5-bisphosphate (PI(4,5)P_2_) into the secondary messengers diacylglycerol (DAG) and Inositol trisphosphate (IP_3_) which in turn activate protein kinase C (PKC) and liberate Ca^2+^ from internal stores. PKC phosphorylation of Ca^2+^ channels, (M)Unc18, SNAP25 and Synaptotagmin modulates neurotransmitter release^5, 30, 31, 32, 33^ and genetic experiments in *C. elegans* suggested a connection to (M)Unc13 as well^34^. However, whether neuromodulators potentiate neurotransmitter release via direct action of the signaling lipids DAG and PI(4,5)P_2_ on (M)Unc13 and how this might affect the protein at the synapse remains to be established.

In insects, octopamine, the invertebrate homolog of mammalian noradrenaline, is a major neuromodulator of synaptic structure and function, locomotion, food seeking and reproduction behavior, as well as learning and memory^23, 35, 36, 37, 38, 39, 40, 41^. At the *Drosophila melanogaster* neuromuscular junction (NMJ), acute octopamine application enhances synaptic transmission, elevates cAMP levels, and triggers the release of neuropeptides, while prolonged exposure enhances neural growth^23, 42, 43^.

Octopamine can also signal via the Gαq/PLC pathway^44, 45^, and here we investigate its role in fast synaptic potentiation in connection to the neurotransmitter release machinery. We combine electrophysiological recordings, GCaMP8-based presynaptic Ca^2+^ influx imaging, confocal microscopy analyses, and live single-molecule imaging at the *Drosophila* larval NMJ with genetic and pharmacological manipulations. We also establish the organismal relevance by behavioral profiling. Our findings point to a requirement of presynaptic OAMB octopamine receptors, PLC and Unc13 for fast potentiation. We provide a mechanism of how this monoamine signaling engages the Unc13 C1 domain for fast recruitment into subsynaptic nanoclusters to enhance synaptic output on a minute’s timescale

## Results

To identify molecular mechanisms of how neuromodulators rapidly modulate synaptic transmission we investigated the molecular pathway for acute octopamine signaling at the *Drosophila melanogaster* 3^rd^ instar larval NMJ.

### Octopamine treatment increases neurotransmitter release and enhances BRP and Unc13 signals within 1 minute

We first tested whether acute octopamine treatment affected synaptic transmission in current-clamp recordings from muscle 6 NMJs. We quantified miniature excitatory postsynaptic potentials (mEPSPs), reporting on the spontaneous, isolated fusion events of individual neurotransmitter-containing SVs; and evoked excitatory postsynaptic potentials (eEPSPs), reporting on AP-evoked, synchronized SV fusion. The Ca^2+^ concentration in the external solution was 0.4 mM, which is sub physiological (∼1.5 mM) but typical for current clamp experiments at the NMJ to prevent muscle contraction and to ensure near-linear summation of mEPSPs to eEPSPs^46^. Exposing larval filets to 20 µM octopamine for 1 minute neither affected the amplitude nor the frequency of mEPSPs (Fig. 1A; S1C) but significantly potentiated the amplitude of AP-induced eEPSPs (Fig. 1A, Fig. S1A), consistent with previous reports^23^. We confirmed that potentiation depended on octopamine and not the simulation protocol in separate experiments without treatment (Fig. S1A). This indicates that octopamine increases the quantal content, the number of neurotransmitter-containing vesicles fusing in response to an AP.

**Figure 1:**
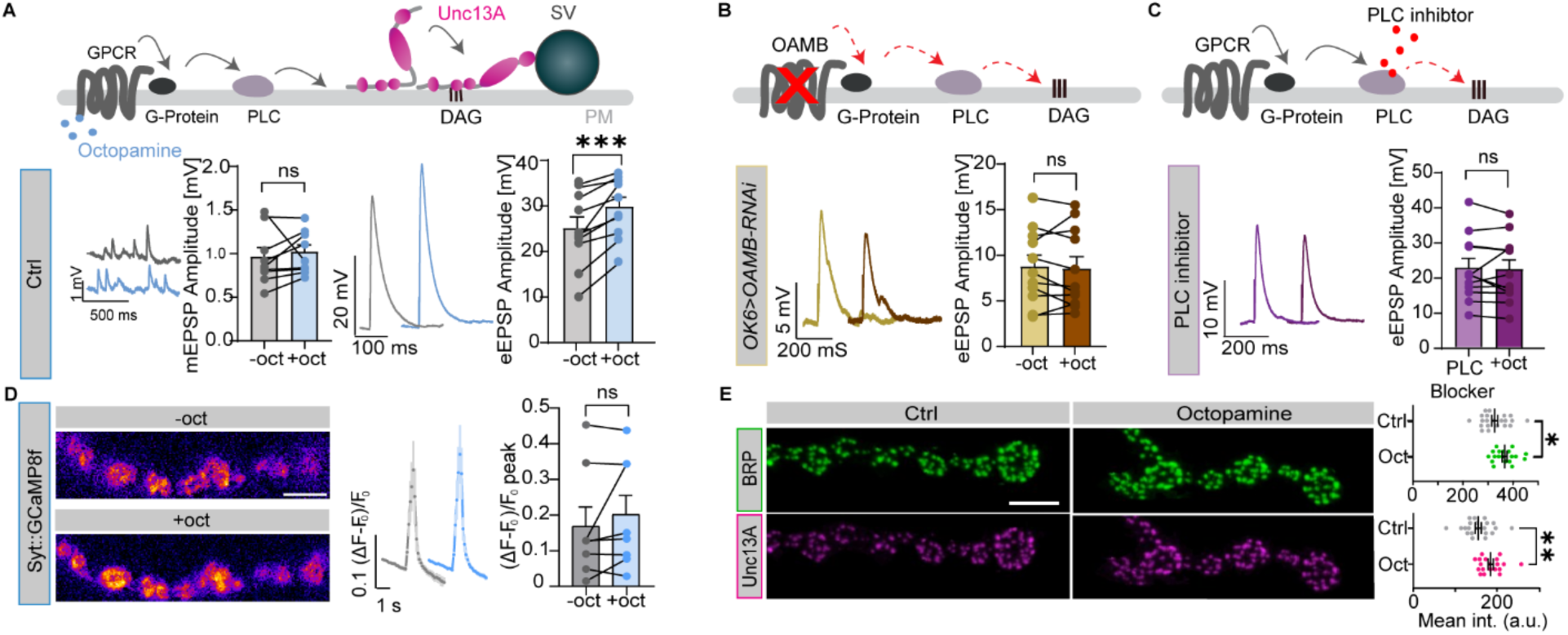
1-minute octopamine treatment potentiates neurotransmitter release via its receptor, does not grossly affect presynaptic Ca^2+^ influx, rapidly enhances BRP and Unc13A AZ signals. (**A**, **B** and **C**) A schematic overview of the octopamine-induced synaptic potentiation pathway, highlighting the involvement of GPCRs, G-proteins, PLC, DAG, and Unc13A, with indications of OAMB knockdown (**B**) and PLC blockage (**C**) with analysis of current clamp recordings (muscle 6 NMJs, 0.4 mM extracellular Ca^2+^) of spontaneous (left) and AP-evoked synaptic activity (right). (**A**) Left: representative example traces of spontaneous mEPSPs from control (Ctrl) animals (wildtype, *w1118*) before (grey, top) and 1-minute after 20 µM octopamine incubation (blue, bottom) together with a cell-wise quantification of mEPSP amplitudes before (-oct, grey) and 1 minute after incubation with 20 µM octopamine (+oct, blue). Right: representative AP-evoked eEPSP responses (average of five repetitions in one cell) from control synapses before (grey) and after octopamine incubation (blue) and quantification of eEPSP amplitudes (-oct, grey; +oct, blue). (**B**) Right: representative AP-evoked eEPSP responses (average of five repetitions in one cell) OAMB KD animals before 1-minute octopamine treatment (yellow) followed by 1-minute incubation with 20 µM octopamine (+oct, brown). (**C**) Representative AP-evoked eEPSP responses (average of five repetitions in one cell) in Ctrl animals (w1118) with 1-minute PLC inhibitor incubation (U73122, 1 µM; light purple) and followed by 1-minute incubation with 20 µM octopamine (dark purple) with quantification of eEPSP amplitudes. (**D**) Left and center: representative images and quantification (averages and SEMs of each recording frame) of GCaMP8f fluorescence signals from muscle 4 NMJs, before (grey, -oct) and after (blue, +oct) octopamine 1-minute incubation. Right: paired comparison of the peak fluorescence signal before and after octopamine incubation. (**E**) Left: Representative confocal images of muscle 4 NMJs of segment A3-A5 from third instar wild type (Ctrl, w1118) *Drosophila* larvae in which larvae are labeled with antibodies labelling BRP (green, top) and Unc13A N-term (pink, bottom) fluorescence signals in either 1-minute DMSO or 1-minute octopamine incubation test conditions. Right: Quantification of averaged NMJ BRP and Unc13A puncta signal in DMSO or octopamine conditions in w1118 larvae. Number of cells (n) and animals (N) investigated: n/N(*Ctrl*, **A**) = 12/12; n/N(OAMB KD, **B**) = 11/11; n/N(PLC inhibitor, **C**) = 12/12; n/N(Syt::GCaMP8f, **D**) = 8/8; n/N(DMSO, **E**) = 21/7, n/N(octopamine, **E**) = 19/7. For exact genotypes see methods. Data depict mean values ± SEM. Statistical analysis with paired parametric t-tests (**A, B** and **D**), unpaired, Mann-Whitney U test (**C**), or unpaired t-test (**E**). n.s., p > 0.05; *p % 0.05; **p % 0.01; ***p % 0.001. Scale bars: 5 µm (**D**) and (**E**)

To investigate the signaling pathway, we tested for the involvement of octopamine receptor in fast synaptic potentiation. While the receptors Octβ1R and Octβ2R induce long-lasting structural changes at the NMJ and are essential for learning^23, 24, 47, 48, 49^, the role of the OAMB receptor -which is also involved in learning and expressed in the motoneurons innervating the NMJs studied here^45, 50, 51^-has thus far not been explored in the context of fast potentiation. We therefore investigated the physiological consequences of its cell specific knockdown in motoneurons. We found no obvious effect of presynaptic OAMB receptor knockdown (OAMB KD, *OK6-Gal4>UAS-OAMB-RNAi*) on AP-evoked baseline synaptic transmission (Fig. S1F). However, acute octopamine-induced potentiation of AP-evoked transmission was completely blocked (Fig. 1B, S1H) (but present in the respective control animals tested in parallel, Fig. S1G) pointing to a requirement of presynaptic OAMB receptors for this rapid potentiation.

Since OAMB receptors can signal via the Gαq pathway^44, 52^, we next examined whether octopamine-induced potentiation depended on PLC both by acute pharmacological inhibition (Fig. 1C) and its presynaptic knockdown (Fig. S1.2G). In the presence of the PLC inhibitor U73455 (Fig. 1C), octopamine-induced potentiation of eEPSPs was blocked (Fig. 1C), while robust potentiation was seen in parallel recordings using a control drug (U73343) (Fig. S1.2B). Further comparisons between control drug and PLC inhibitor conditions revealed no detectable effects on mEPSP amplitudes and frequencies, or baseline eEPSP amplitudes by these compounds prior to octopamine treatment (Fig. S1.2). To specifically investigate PLC requirement in motor neurons, we knocked down PLC expression presynaptically by targeted RNAi expression (*OK6-Gal4>UAS-PLC-RNAi*). While 1-minute octopamine treatment robustly potentiated eEPSPs in control animals (expressing UAS-PLC-RNAi, but not the OK6-Gal4 driver) (Fig. S1.2E), only a slight eEPSP potentiation was seen following presynaptic PLC knockdown (Fig. S1.2G). Notably, RNAi expression significantly reduced the octopamine-induced potentiation of eEPSP amplitudes compared to controls (Fig. S1.2H). No effect on baseline eEPSP amplitudes was seen, but PLC knockdown increased mEPSP amplitudes compared to the control genotype (Fig. 1.2L). Collectively, these results indicate that the presynaptic Gαq pathway mediates octopamine-induced potentiation.

**Figure 2:**
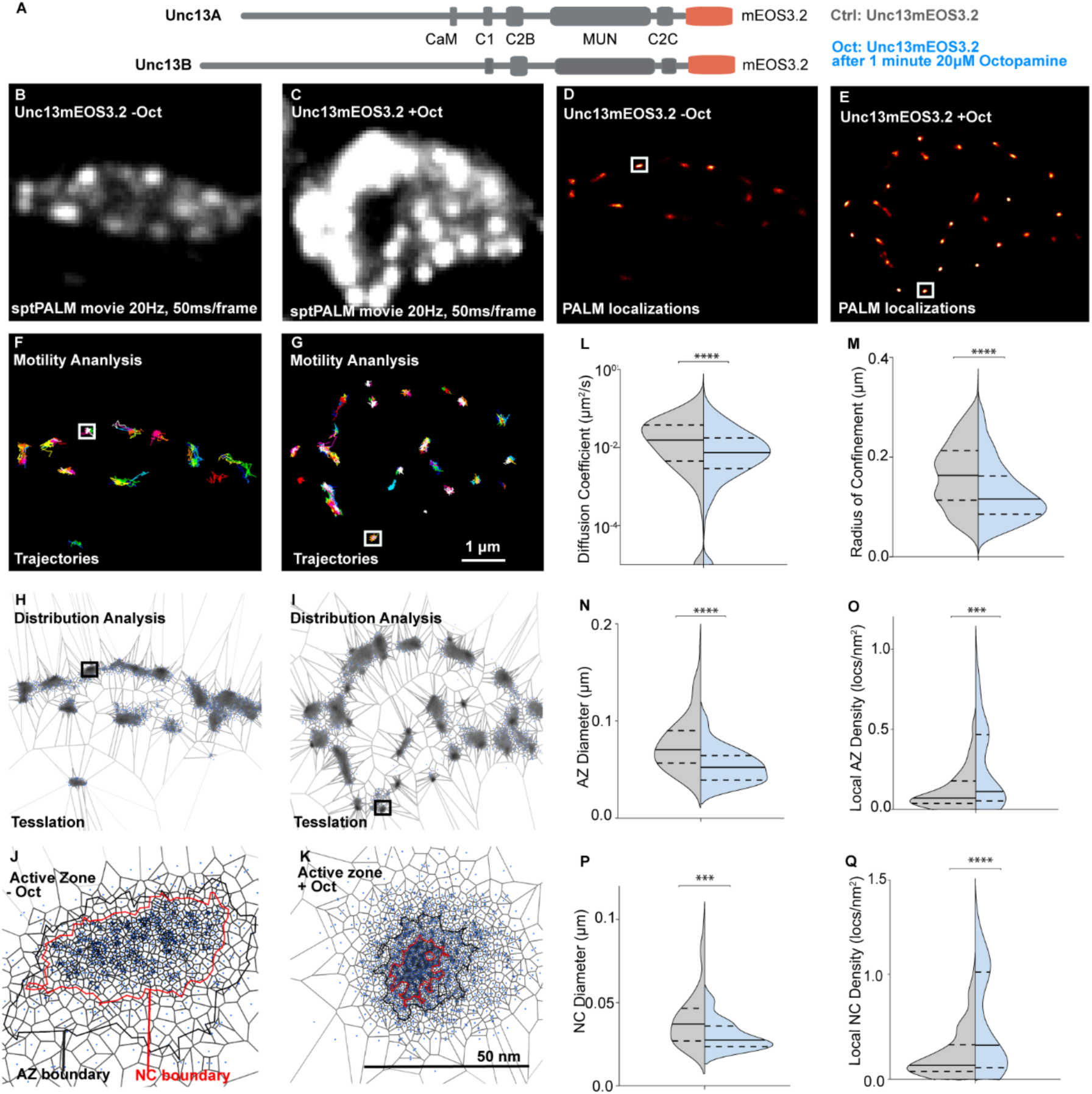
Live *in vivo* imaging of endogenously tagged Unc13 molecules undergo compaction upon acute octopamine treatment. Live sptPALM imaging of Unc13mEOS3.2 at muscle 4 NMJs, performed in 0.4 mM Ca^2+^ and 10 mM Mg^2+^ containing HL3.1, before (Internal Control/Ctrl) and after a 1-minute incubation of 20 μM octopamine in HL3.1 (Oct). (A) Scheme of Unc13A and Unc13B protein showing the regulatory Ca^2+^/calmodulin binding (CaM) domain (only in Unc13A), the DAG/PMA binding C1 domain, the Ca^2+^/phosphoinositide binding C2B domain, the MUN domain, the C2C domain, and an endogenously encoding C-terminal mOES3.2 tag. Images show representative sptPALM recordings (**B** and **C**), PALM localization map (**D** and **E**), trajectory maps (**F** and **G**), and tessellation analysis representations of Unc13mEOS3.2 before and after 1-minute octopamine treatment (**H**-**K**). (**L** and **M**) Quantification of diffusion coefficients and radii of confinement from live Unc13 channel sptPALM imaging. Ctrl NMJs = 2051 single-molecule trajectories; 1-minute octopamine treatment = 2505 single-molecule trajectories. (L-O) Tessellation analysis from the same sptPALM dataset was analyzed in (**B**-**M**) for diameters and densities of Unc13 localizations within AZ (**N**-**O**) and NC cluster (**P**-**Q**) boundaries. Here, 7 animals per condition generated 106 AZs (control) and 86 AZs (Oct), respectively, that were analyzed. Number of NMJs (n), animals (N), number of AZs (X) and number of individual trajectories (Y) investigated: n/N/X/Y(*Ctrl*) = *7*/7/106/2051; n/N X/Y (Oct) = 7/7/86/2505; Data depict mean values ± SEM. Statistical significance is denoted as asterisks: **P < 0.01, ***P < 0.001, and ****P < 0.0001. Data distribution was statistically tested with a Kolmogorov-Smirnov test. AU, arbitrary units. Scale bars, 1 μm (**B**-**G**) and 50 nm (**H** and **I**).

Octopamine can increase Ca^2+^ influx through voltage gated channels^53^ which could explain the observed potentiation. We therefore investigated whether enhanced AP-induced Ca^2+^ influx was seen upon octopamine treatment in live imaging experiments using the fluorescent Ca^2+^ indicator Synaptotagmin::GCaMP8f selectively targeted to the presynaptic active zone (AZ)^54^. We quantified baseline-corrected and normalized fluorescence intensity changes induced by stimulating muscle 6 NMJs with 20 APs at a frequency of 20 Hz in a similar 0.4 mM [Ca^2+^] external solution (Fig. 1D). To evaluate whether octopamine enhanced AP- induced Ca^2+^ influx, the same NMJ was stimulated twice, before and 1 minute after incubation with 20 µM octopamine. This resulted in a very similar presynaptic Ca^2+^ response and no statistically significant changes were seen (Fig. 1D). The repeated stimulation and imaging alone (without octopamine) also revealed comparable Ca^2+^ responses to both stimuli (Fig. S1D). While we cannot exclude relevant differences in Ca^2+^ influx below the detection threshold of this method, these findings suggest that under the conditions investigated here, octopamine is unlikely to dramatically alter AP-induced Ca^2+^ influx.

Increasing the number of neurotransmitter release sites also enhances the quantal content. Unc13 is a limiting component of these sites and recruited to the AZ by the scaffolding protein Bruchpilot (BRP)^55, 56, 57^. We therefore tested whether acute octopamine treatment affected their signals at NMJs fixed and stained using antibodies against BRP and Unc13A and imaged by confocal microscopy. Comparing the mean synaptic fluorescence intensity values of BRP and Unc13A between animals with or without octopamine treatment for 1 minute prior to tissue fixation revealed an increase of signals for both proteins (Fig. 1E). While this finding qualitatively aligns with previous observations during presynaptic homeostatic potentiation (PHP) on a timescale of 10 minutes^7, 58^, these data demonstrate an even faster (1-minute) adaptation of the presynaptic AZ composition upon octopamine-induced potentiation.

### Rapid nanoscale enrichment of Unc13 with octopamine treatment

Recently, a compaction of Unc13 immunolabelled molecules was reported using single molecule imaging at the fly NMJ in response to 10 minutes of treatment inducing homeostatic potentiation in fixed samples^8^. We therefore also performed single molecule imaging but chose a live imaging approach combined with a CRISPR Cas9 mediated tagging of the endogenous Unc13 to directly assess the protein’s level and localizations without the need of antibody labelling. For this purpose, an mEOS3.2 tag was fused to the wildtype Unc13 gene (Fig. 2A) similar to recent analyses for BRP and the calcium channel cacophony^11, 56^. We confirmed that this modification did not affect basal synaptic properties (Fig. S2.1). We wanted to capture the rapid adaptation of the nanoscopic Unc13 distribution at the synapse during the fast potentiation induced by octopamine and therefore developed an assay to combine acute pharmacological application with live single molecule imaging of endogenously tagged Unc13. We performed live sptPALM imaging of Unc13mEOS3.2 populations at a Ib type muscle 4 NMJs before drug application as an internal control. Following this, a new muscle 4, Ib, NMJ in the same animal was identified, 20 μM octopamine applied and the NMJ immediately imaged, starting 1 minute after drug application for a duration of 8.5 minutes at an acquisition rate of 20 Hz (Fig. 2, further details described in the methods section). This allowed us to capture the behavior of individual Unc13 molecules at NMJs of the same animal before and immediately after octopamine application.

We first analyzed the Unc13 molecule dynamics. Notably, octopamine treatment reduced Unc13 motility, evident in a profound decrease of median diffusion coefficient (median diffusion coefficient: internal controls: 0.0155 μm^2^/s; octopamine treated: 0.00774 μm^2^/s) (Fig. 2I). In addition, the mean radius of confinement of Unc13 trajectories significantly decreased by 30% after 1-minute octopamine treatment compared to the synapses of paired internal controls (internal control, 163.2 nm; octopamine treated, 115.9 nm) (Fig. 2J). Parallel experiments without treatment established that the repeated imaging itself did not affect this (Fig. S2.2). Thus, octopamine treatment promotes immobilization and confines the spread of Unc13 molecules at the AZ within a few minutes.

We used an SR tessellation analysis to calculate the nanoscale distribution of Unc13 molecules within the broader AZ cluster and the central nanocluster from PALM localization data (Fig. 2N-O and P-Q) by calculating the diameters and densities occupied by Unc13 localizations. Signals of Unc13 localizations underwent a significant reduction in AZ cluster diameters (by about 25.8%; Fig. 2N). Meanwhile, AZ Unc13 localization densities significantly increased by 143.5% within 1 minute of octopamine application compared to synapses before treatment (Fig. 2O). Similarly, within nanocluster boundaries a significant Unc13 density increase (about 96.5%) was observed, and the nanocluster diameters decreased significantly (by about 25.9%; Fig. 2P and Q). These results point to a rapid compaction to high-density nanoclusters rather than an absolute increase in Unc13 protein numbers mediating the fast synaptic potentiation seen upon octopamine application.

To investigate whether the observed increase in protein density was partly due to an increase in Unc13 molecules at the AZ, we analyzed a subset of the same imaging data in Figure 2 data to quantify the number of stepwise transitions during bleaching of Unc13mEOS3.2 fluorophore. This revealed 8-17 Unc13 molecules per AZ (Fig. S2.3) at rest. The number of Unc13 molecules estimated is based on the fluorescent signal of mEOS and biased by the blinking emission properties of the fluorophore (mEOS3) and the unknown dark state of the fluorophore within the cellular environment. Thus, the counting potentially underestimates the real number of Unc13 molecules per AZ. Despite these uncertainties, this approach allows a comparison of relative numbers across imaging conditions. This revealed similar counts for Unc13 molecules before and after octopamine-induced potentiation (Fig. S2.3). Thus, our results point to a rapid compaction to high-density nanoclusters rather than an absolute increase in Unc13 protein numbers mediating the fast synaptic potentiation seen upon octopamine application.

### Octopamine-induced potentiation depends on presynaptic Unc13A

Both *Drosophila melanogaster* splice isoforms Unc13A and -B are present at the 3^rd^ instar NMJ (Fig. 3A) and both isoforms are tagged in the experiments above. However, neurotransmitter release at this synapse almost exclusively depends on Unc13A^57^. To explore whether Unc13A plays a direct role in this rapid potentiation, we tested whether potentiation was disrupted upon Unc13A knockdown (KD) in motor neurons (using OK6-Gal4)^7, 59^. Compared to control animals (expressing only the UAS-Unc13A-RNAi), presynaptic Unc13A knockdown (Unc13A KD, *OK6-Gal4>UAS-Unc13A-RNAi*) increased mEPSP amplitudes, while no changes were observed in frequencies, and decreased eEPSP amplitudes (Fig. 3C, S3.1) in qualitative agreement to the Unc13A null mutation^57^. While neurotransmitter release was potentiated at NMJs from control animals upon 1-minute 20 µM octopamine treatment ((Fig. 3B) similar as in wildtype animals, Fig. 1A), no potentiation was seen upon presynaptic Unc13A KD (Fig. 3D). This points to a direct role of Unc13A in mediating the fast neurotransmitter potentiation by octopamine.

**Figure 3:**
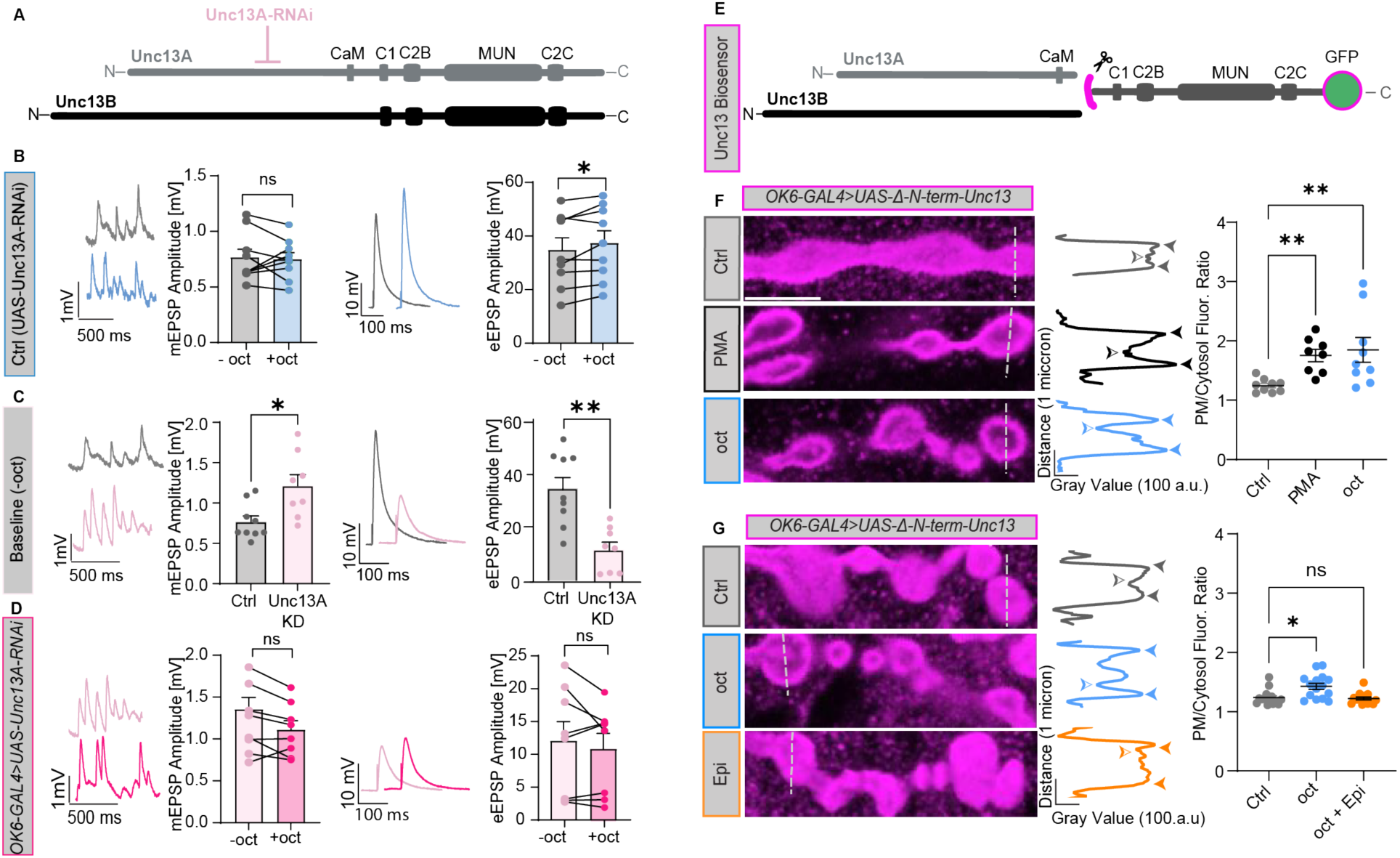
Octopamine induced synaptic potentiation requires Unc13A and a C terminal Unc13-fragment is recruited to the plasma membrane by the phorbol ester PMA and octopamine. (**A**) Scheme of Unc13A and -B isoforms and the RNAi approach for Unc13A knockdown. Different domains are shown: regulatory Ca^2+^/calmodulin binding (CaM) domain (Unc13A specific), DAG/PMA binding C1 domain, Ca^2+^/phosphoinositide binding C2B domain, the functionally essential MUN domain, and the C2C domain linked to SV association. (**B**, **C** and **D**) Analysis of current clamp recordings (muscle 6 NMJs) of AP-evoked synaptic activity recorded in the presence of 0.4 mM extracellular Ca^2+^. (**B**) Left: Representative example traces of spontaneous mEPSPs from control animals (ctrl, UAS-Unc13A-RNAi) together with a cell-wise quantification of mEPSP amplitudes before (-oct, grey) and 1 min after incubation with 20 µM octopamine (+oct, blue). Right: Representative AP-evoked eEPSP responses (average of five repetitions in one cell) and cell-wise comparison of eEPSP amplitudes in control synapses before (-oct, grey) and after octopamine incubation (+oct, blue). (**C**) Left: representative example traces of spontaneous mEPSPs and cellwise quantification of mEPSP amplitudes from Ctrl and Unc13A knock down (Unc13A KD) synapses. Right: representative AP-evoked eEPSP responses (average of five repetitions in one cell) from Ctrl and Unc13A KD synapses and quantification of eEPSP amplitude. (**D**) Left: representative example traces of spontaneous mEPSPs from Unc13A KD synapses (Unc13A KD, *OK6-Gal4>UAS-Unc13A-RNAi*) before (light pink, top) and 1-minute after 20 µM octopamine incubation (pink, bottom) together with a cell-wise quantification of mEPSP amplitudes before (-oct, light pink) and 1 min after incubation with 20 µM octopamine (+oct, pink). Right: representative AP-evoked eEPSP responses (average of five repetitions in one cell) from control synapses before (light pink) and after octopamine incubation (pink) and quantification of eEPSP amplitudes (-oct, light pink; +oct, pink). (**E**) Scheme of the Unc13 A and B fragment which it is truncated before the C1 domain and contains a C-terminally fused GFP. (**F**) Left: Representative confocal images of muscle 4 NMJs of segment A3–5 from Unc13 Biosensor (*OK6-Gal4>UAS-Δ-N-term-Unc13-GFP*) third-instar *Drosophila* larvae incubated with Ctrl (DMSO, grey) for 1 minute as negative control, with 2 µM PMA in DMSO for 10 minutes (black) as a positive control, and 20 µM octopamine in DMSO for 1 minute (blue). Intensity profiles were read out at the locations indicated by the dashed lines. Middle: Line profiles from respective images representing Unc13A-GFP fluorescent intensities. The max intensity values in the line profiles close to the bouton edge (plasma membrane; solid arrows) were averaged and then divided by the minimum fluorescence intensity in the cytosol (hollow arrow) to generate a plasma membrane/cytosol fluorescence ratio of each bouton. All bouton ratios per NMJ were then averaged and compared across animals in the graphs on the right. Right: NMJ ratios across animals. (**G**) Performed same as in **B**, where Unc13 Biosensor (*OK6-Gal4>UAS-Δ-N-term-Unc13- GFP*) 3^rd^ instar larvae were incubated for 1 minute in 20 µM octopamine in DMSO alone (blue), or after pre-treatment for 5 minutes in 1 µM epinastine in DMSO (orange). Number of cells (n) and animals (N) investigated: n/N(Genetic ctrl) = 9/9; n/N(Unc13A KD) = 8/8; n/N(Ctrl, **F**) = 9/3; n/N(PMA, **F**) = 8/3.; n/N(oct, **F**) = 9/3; n/N(Ctrl, **G**) = 15/5; n/N(oct, **G**) = 15/5; n/N(oct + Epi, **G**) = = 15/5. For exact genotypes see methods. Data depict mean values ± SEM. Statistical analysis with unpaired, Mann-Whitney U test (**F**), or Kruskal-Wallis Test (**G**), or with paired parametric t-tests (**B, C** and **D**). n.s., p > 0.05; *p % 0.05; ***p % 0.001. Scale bar in (**F**): 5 µm.

### Octopamine signaling recruits a delocalized Unc13 C-terminal fragment to synaptic plasma membranes

Because Unc13A levels correlate with synaptic strength and BRP recruits Unc13A via interactions with the Unc13A N-terminus^55, 56, 57^, we wondered to what extent potentiation was secondary to increased BRP levels, or whether octopamine engaged Unc13 directly. We therefore uncoupled its interaction with BRP by deleting the Unc13 N-terminus in a GFP-tagged C-terminal fragment overexpressed in motoneurons (using the OK6-Gal4 driver) to investigate whether 1-minute application of octopamine (20 µM) resulted in any detectable change in protein localization. This construct was delocalized at rest as seen in confocal images (Fig. 3E)^56^ but still harbors the Unc13 lipid binding C1 and C2B domains. A similar fragment of the mammalian ortholog Munc-13 reacts to phorbol ester treatment by rapid translocation from the cytosol to the plasma membrane in HEK cells^60^ and we confirmed that this was also the case for our construct at the NMJ treated with 2 µM PMA (Fig. 3F). We quantified the distribution of this fragment in single confocal slices using line profiles across NMJ boutons (Fig. 3F, G; dotted lines). This revealed rather homogenous intensity levels of the GFP fluorescence in control (DMSO-treated) NMJs, while treatment with the phorbol ester PMA (2 µM for 10 minutes) enhanced the fluorescence intensity at the edges of the bouton while decreasing the intensity in the center of the bouton (Fig. 3F), indicating the fragment’s translocation to the plasma membrane. To quantitatively compare this distribution across animals, we compared the ratios between the highest intensity peaks at the bouton edge with the lowest fluorescent intensity in the bouton center across treatment conditions. This revealed that 1-minute exposure to 20 µM octopamine similarly redistributed the Unc13 C-terminal fragment to the plasma membrane as PMA (Fig. 3F). To check whether this signaling depended on octopaminergic GPCRs, we used epinastine to specifically block them^61^. Indeed, pre-incubation of larval fillets with epinastine (1 µM for 5 minutes) fully blocked the octopamine-induced translocation of the Unc13 C-terminal fragment (Fig. 3G). Our data indicate that octopamine can recruit the C-terminal portion of Unc13 to the plasma membrane. These findings indicate a responsiveness by Unc13 encoded in its C-terminal domains which we next investigated further.

### Mutations in the Unc13 lipid binding C2B and C1 domains enhance neurotransmitter release, reduce synaptic Unc13A signals and attenuate octopamine-induced potentiation

We wanted to investigate whether Unc13’s C1 and C2B domains which bind to the two signaling lipids of the Gαq pathway (DAG and PI(4,5)P_2_) contribute to octopamine-induced potentiation. Both domains are needed for membrane binding and were shown to exert auto-inhibitory functions^62, 63^. However, cellular pathways that modulate synaptic function via these domains are not known. We generated mutant flies by CRISPR Cas9 gene editing of specific amino acids in the two domains. As both domains are present in either Unc13 splice isoform, this approach targeted both Unc13A and -B. In the C2B domain mutant, a central lysine was mutated to a tryptophane (Unc13-C2B^KW^; K1862W in Unc13A) (Fig. 4A) which in the mammalian ortholog was shown to enhance synaptic transmission in primary hippocampal neurons and to increase the affinity of the C2B domain to bind PI(4,5)P_2_-containing membranes^64^. The mutation in the C1 domain changed a central histidine to a lysine (Unc13-C1^HK^; H1723K in Unc13A) (Fig. 4D) which was shown to enhance synaptic transmission in cultured hippocampal neurons by lowering the energy barrier for vesicle fusion^16, 17, 65^. We first confirmed that these mutations also enhanced neurotransmitter release in flies by performing current clamp recordings at muscle 6 NMJs. This revealed an increase of AP-induced eEPSPs for both mutants when compared to wildtype animals (Fig. 4B and E right). At the same time, no detectable changes of mEPSP amplitudes or frequencies were seen (Fig. 4B and E left; Fig. S4.1), consistent with a chronic presynaptic potentiation. To investigate whether these mutations affected the morphology of the synaptic AZ and/or the AZ levels of Unc13A, confocal analysis was performed following immunostaining against BRP and Unc13A. This revealed reduced mean synaptic fluorescent intensities in the Unc13A channel in both the Unc13-C2B^KW^ and Unc13-C1^HK^ mutants compared to control animals (wildtype, w1118), whereas BRP signals were similar (Fig. 4C and F). The fact that neurotransmitter release at these mutant NMJs was increased, while Unc13A signals were decreased argues for an intrinsically enhanced activity of Unc13 caused by these mutations. The reduced levels might be a result of a secondary, homeostatic compensation in response to the hyperactive effect of the Unc13-C2B^KW^ and Unc13-C1^HK^ mutations (see discussion).

**Figure 4:**
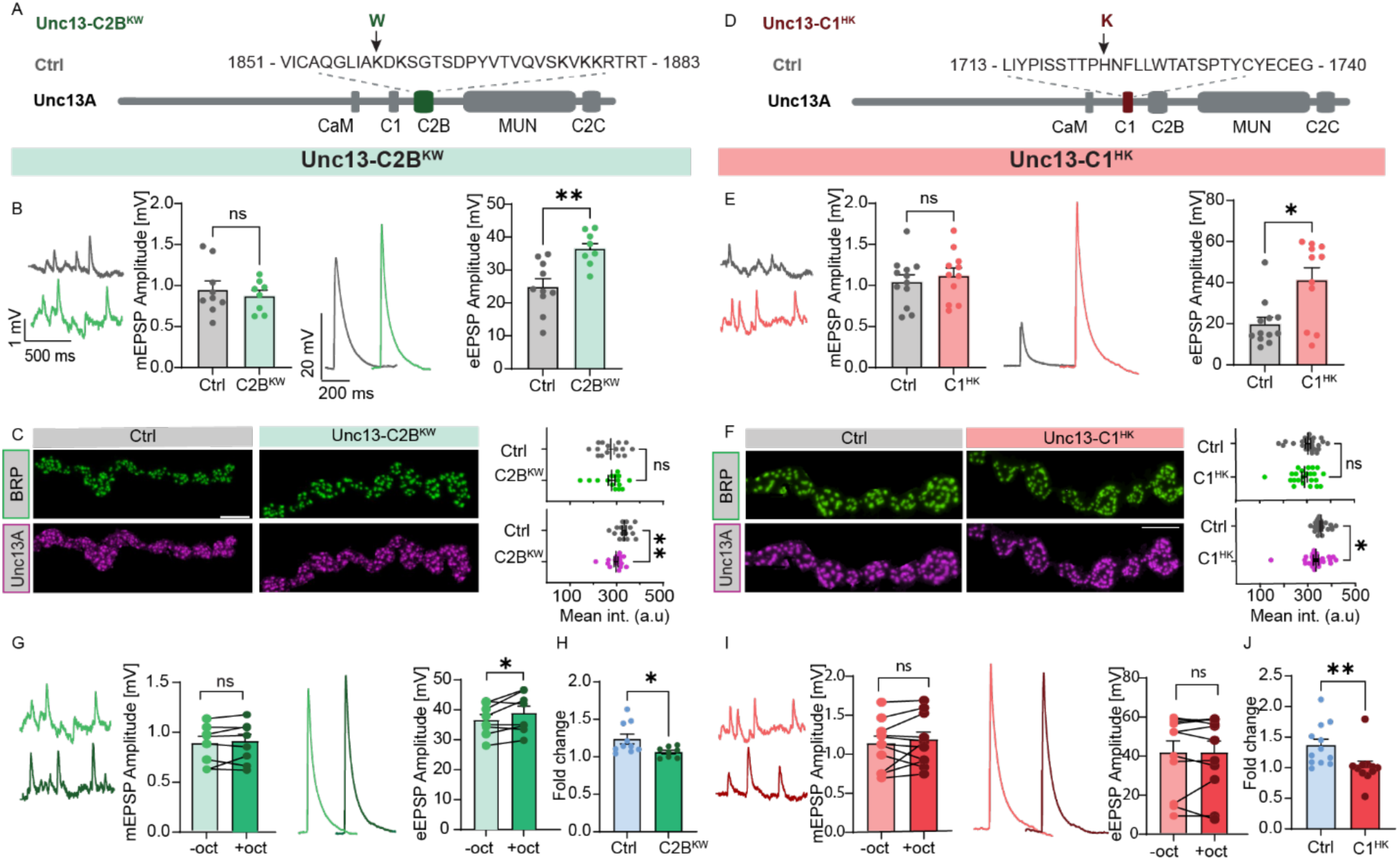
Enhanced neurotransmitter release, reduced Unc13A signals, and diminished octopamine-induced potentiation in Unc13-C1^HK^ and Unc13-C2B^KW^ domain mutants. (**A**, **D**) A schematic representation of Unc13A protein showing the regulatory Ca^2+^/calmodulin binding (CaM) domain, the DAG/PMA binding C1 domain, the Ca^2+^/phosphoinositide binding C2B domain, as well as the MUN and C2C domain. Left: Unc13-C2B^KW^ mutant (K1862W in Unc13A) and right: Unc13-C1^HK^ mutant (H1723K in Unc13A). Arrows indicate the amino acid changes. (**B**, **G**, **E** and **I**) Analysis of current clamp recordings (muscle 6 NMJs, 0.4 mM extracellular Ca^2+^) of spontaneous (left) and AP-evoked synaptic activity (right) in control (Ctrl) and Unc13-C2B^KW^ mutant/Unc13-C1^HK^ animals. (**B**) Left: representative example traces of spontaneous mEPSPs from Ctrl (grey) and Unc13-C2B^KW^ mutant synapses (light green) and cell-wise quantification of mEPSP amplitudes. Right: representative AP-evoked eEPSP responses (average of five repetitions in one cell) from Ctrl (grey) and Unc13-C2B^KW^ mutant synapses (light green) and cell-wise quantification of eEPSP amplitude. (**C**) Left: Representative confocal images of muscle 4 NMJs of segment A3–5 from third-instar *Drosophila* larvae of the indicated genotypes, stained with antibodies against BRP (green, top) and the N-terminal region of Unc13A. (purple, bottom, see methods). Right: Quantification of mean intensities of BRP (top) and Unc13A (bottom) in Ctrl and Unc13-C2B^KW^ mutant NMJs. (**G**) Left: representative example traces of spontaneous mEPSPs and cell-wise quantification of mEPSP amplitudes before octopamine (-oct, light green) and after 1-minute incubation with 20 µM octopamine (+oct, dark green) at Unc13-C2B^KW^ mutant synapses. Right: representative AP-evoked eEPSP responses (average of five repetitions in one cell) before octopamine (-oct, light green) and after 1-minute incubation with 20 µM octopamine (+oct, dark green) and quantification of eEPSP amplitude. (**H**) Comparison of the fold change in eEPSP amplitudes after and before 1 minute of octopamine incubation between Ctrl animals (blue) and Unc13-C2B^KW^ (dark green) mutants. (**E**, **F**, **I** and **J**) Same analysis as in (**B**, **C**, **G** and **H**) for Unc13- C1^HK^ mutant. Number of cells (n) and animals (N) investigated: n/N: n/N(*Ctrl,* B, G, H) *=* 10/10 (eEPSP) and 9/9 (mEPSP), n/N(*Unc13-C2B^KW^*, **B**,**G** and **H**) = 8/8(eEPSP) and 8/8 (mEPSP), n/N(*Ctrl,* E, **I**, **J**) = 12/12 (eEPSP) and 12/12 (mEPSP), n/N(*Unc13-C1^HK^*,**E**,**I** and **J**) = 11/11, n/N(Ctrl, **C**) = 15/5; n/N(Unc13- C2B^KW^, **C**) = 15/5, n/N(Ctrl, **F**) = 24/8, n/N(Unc13-C1^HK^, **F**) = 23/8. For exact genotypes see methods. Data depict mean values ± SEM. Statistical analysis with unpaired, Mann-Whitney U test (**B**, **C**, **E**, **F**, **H** and **J**) or with paired parametric t-tests (**G** and **I**). n.s., p > 0.05; *p % 0.05; **p % 0.01. Scale bar in (**C**, **F**): 5 µm.

To probe for a possible engagement of the Unc13 C2B and C1 domain in octopamine-induced potentiation, we next investigated the reactivity of the Unc13-C2B^KW^ and Unc13-C1^HK^ mutants to 1-minute 20 µM octopamine incubation. While octopamine still potentiated neurotransmitter release in animals expressing the Unc13-C2B^KW^ mutation (Fig. 4G), potentiation was lost in Unc13-C1^HK^ mutants (Fig. 4I), consistent with a more direct involvement of the C1 domain in the octopamine-induces potentiation of neurotransmitter release. We therefore investigated the functional consequences of this mutation further.

### Mutation of the Unc13 C1 domain blocks octopamine-induced nanoscale compaction and reduces Unc13 copy numbers

To test if acute octopamine potentiation is in fact dependent on the acute nanoscale compaction of Unc13 as observed in Figure 2, we performed sptPALM imaging on endogenously tagged Unc13-C1^HK^ mutants (Unc13-C1^HK^-mEOS3.2 mutant). We observed that the Unc13-C1^HK^-mEOS3.2 mutant no longer underwent compaction or motility changes upon acute octopamine treatment (Fig. 5L-U). The simultaneous block of potentiation and structural adaptation in these mutants therefore implies that the observed rapid changes in Unc13 mobility and localization seen for the wildtype protein with octopamine are indeed causally linked to the increased neurotransmitter release.

**Figure 5:**
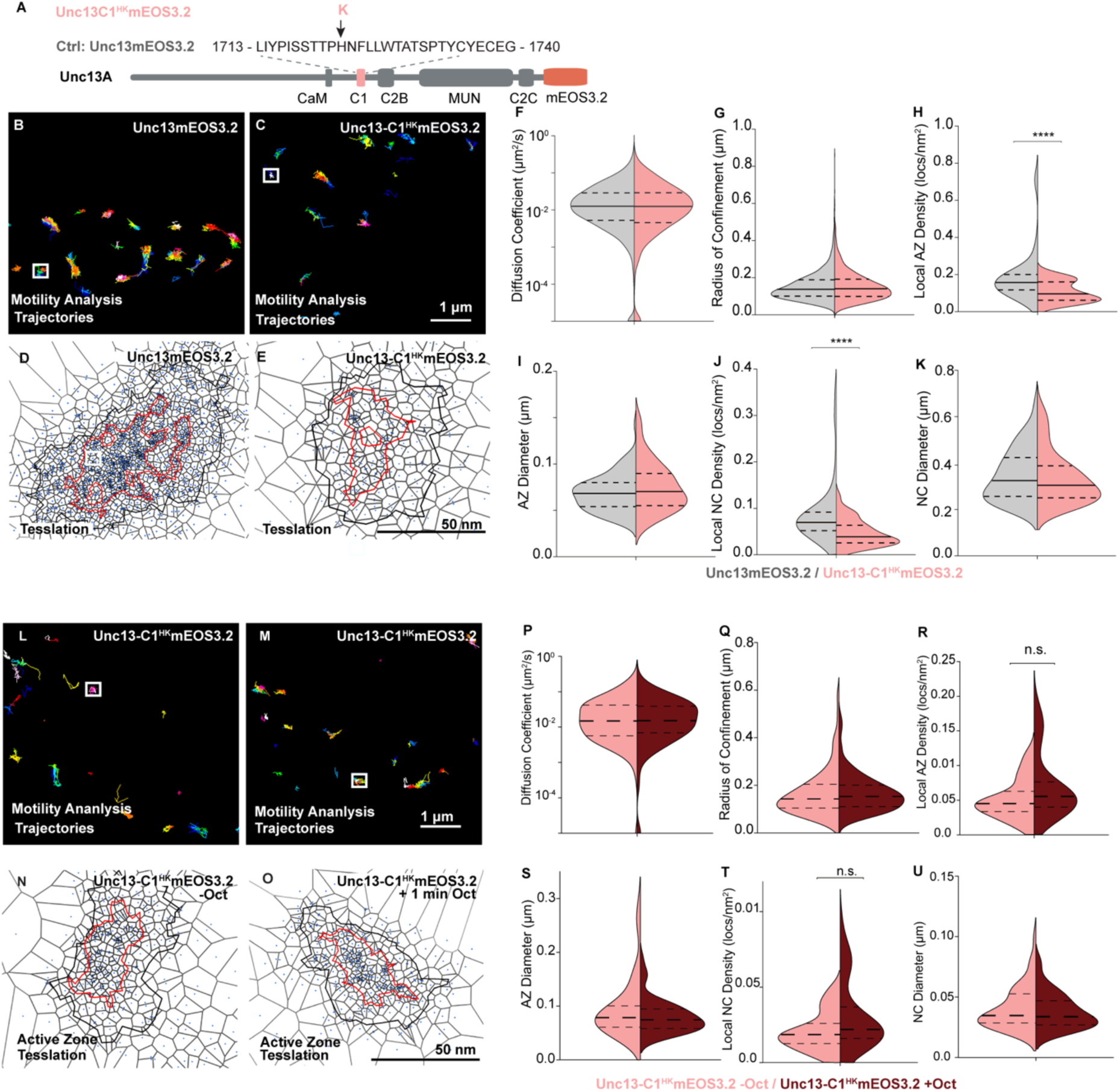
Live *in vivo* imaging of Unc13-C1^HK^ -mEOS reveals reduced Unc13 signals and loss of octopamine-induced Unc13 compaction at AZs. **(A)** Scheme of Unc13A protein showing the regulatory Ca^2+^/calmodulin binding (CaM) domain (only in Unc13A), the DAG/PMA binding C1 domain, the Ca^2+^/phosphoinositide binding C2B domain, the MUN domain, the C2C domain, and an endogenously encoding C-terminal mOES3.2 tag. Unlike in mammals and *C. elegans*, the N-terminal C2A domain is absent in *Drosophila melanogaster*. Unc13-C1^HK^ mutant. Arrow indicates the amino acid change in the C1 domain. (**B**-**E**) Example workflow of sptPALM imaging of endogenously tagged Unc13mEOS3.2 (Ctrl) and Unc13-C1^HK^mEOS3.2 NMJs at muscle 4 NMJs, performed in 1.5 mM Ca^2+^ and 1.9 mM Mg^2+^ containing HL3.1. Images show representative sptPALM recordings movies (generated trajectory maps (**B** and **C**) and SR tessellation map of individual AZs for distribution analysis (**D** and **E**). Analysis from the data in (**B**-**E**) from trajectory maps motility analysis (**F**-**G**) and channel number analysis (Fig. S5.1) and SR tessellation map for distribution analysis (**H**-**K**). (**F** and **G**) Quantification of diffusion coefficients and radii of confinement from live Unc13 channel sptPALM imaging. Control NMJs = 5832 single-molecule trajectories; Unc13-C1^HK^ mutant = 2306 single-molecule trajectories. (**H** - **K**) SR Tessellation analysis performed on the same sptPALM dataset analyzed in (**K** and **L**) extracting diameters and densities of Unc13 localizations in Ctrl and Unc13-C1^HK^ mutant within AZ (**H** and **I**), NC cluster (**J** and **K**) boundaries. A total of 108 AZs (Ctrl) and 111 AZs (Unc13-C1^HK^ mutant) were analyzed from six animals for each condition. Number of NMJs (n), animals (N), number of AZs (X) and number of individual trajectories(**Y**) investigated: n/N/X/Y (Unc13-C1^HK^ mutant -Oct) = 10/6/108/5832; n/N (Unc13-C1^HK^ mutant +Oct) = 10/6/111/2306. Images in (**L**-**O)** represent sptPALM recordings, trajectory maps (**L** and **M**), and AZ tessellation analysis (**N** and **O**) representations of Unc13- C1^HK^mEOS3.2 before and after a 1-minute octopamine (Oct) treatment. (**J** and **K**) Quantification of diffusion coefficients and radii of confinement from live Unc13 channel sptPALM imaging. Control NMJs: = 264 single-molecule trajectories; Unc13-C1^HK^ mutant -Oct mutant = 183 single-molecule trajectories; Unc13C1^HK^ mutant +Oct. (**R**-**U**) Tessellation analysis from the same sptPALM dataset was analyzed in (**L-U**) to (**I**) for diameters and densities of Unc13 in Unc13- C1^HK^mEOS3.2 mutant before /after treatment localizations within AZ (**R**-**S**) and NC cluster (**T**-**U**) boundaries. Here, A total of 45 AZs (Unc13-C1^HK^ mutant -Oct) and 52 AZs (Unc13-C1^HK^ mutant +Oct), were analyzed from six/seven animals per condition. Number of NMJs (n), animals (N), number of AZs (X), and the number of individual trajectories (**Y**) investigated: n/N/X/Y(Unc13- C1^HK^ mutant -Oct) = 6/6/45/264; n/N/X/Y (Unc13-C1^HK^ mutant +Oct) = 7/7/52/183. Statistical significance is denoted as asterisks: \*\**P* < 0.01, \*\*\**P* < 0.001, and \*\*\*\**P* < 0.0001. Data distribution was statistically tested with a Kolmogorov-Smirnov test. AU, arbitrary units. Scale bars, 1 μm (**B**, **C**, **L** and **M**) and 50 nm (**D**, **E**, **N** and **O**).

A genotype-specific comparison of Unc13 motility and nanoscale localization revealed no differences between the tagged wildtype protein or the Unc13-C1^HK^ mutant (Fig. 5B-K). However, it presented a significantly reduced protein abundance following Unc13-C1^HK^ mutation, consistent with the lower signals seen in confocal analysis (Fig. 4F). This was further supported by a significant reduction in the number of Unc13-C1^HK^-mEOS3.2 proteins detected by stepwise photobleaching (Fig. S5.1). Together, engagement of the Unc13-C1 domain is needed to mediate rapid nanoscale compaction and its chronic mutation leads to markedly reduced protein copy numbers, potentially due to a homeostatic response to offset the increased intrinsic activity of the Unc13-C1^HK^ mutant protein (see discussion).

### Unc13-C1^HK^ mutation potentiates neurotransmitter release at physiological Ca^2+^ and below, causes short-term depression and blocks phorbol ester induced potentiation

We sought to investigate whether the potentiation caused by the Unc13-C1^HK^ mutation depended on the external Ca^2+^ concentration. Moreover, we wanted to analyze whether it affected short-term plasticity (STP), the change in synaptic transmission upon repetitive stimulation of the synapse on the millisecond timescale. We therefore performed two-electrode voltage clamp (TEVC) experiments, where the membrane potential of the muscle is kept constant by injecting a current. This prevents muscle contractions with more neurotransmitter release at physiological external Ca^2+^ concentrations (∼1.5 mM [Ca^2+^]_ext_) and above or upon repetitive stimulation. The injected currents needed to maintain the membrane potential provide a direct readout of synaptic transmission. TEVC recordings from muscle 6 NMJs were obtained from either wildtype flies (control, Ctrl) or ones bearing the Unc13-C1^HK^ mutation. Excitatory Postsynaptic Currents (EPSCs) were measured in response to paired AP-stimuli (10 ms apart) at successively increased Ca^2+^ concentrations in the external medium (Fig. 6A). Comparison of the amplitude of the first evoked EPSC (eEPSC_1_) revealed larger responses in Unc13-C1^HK^ mutant flies at physiological (1.5 mM) external Ca^2+^ concentrations and below (Fig. 6B). The mutation also affected STP behavior as seen by lower paired-pulse ratios (PPRs) of the eEPSC amplitudes successively induced by repetitive stimulation (PPR = eEPSC_2_/eEPSC_1_) at all investigated Ca^2+^ concentrations (Fig. 6C).

**Figure 6:**
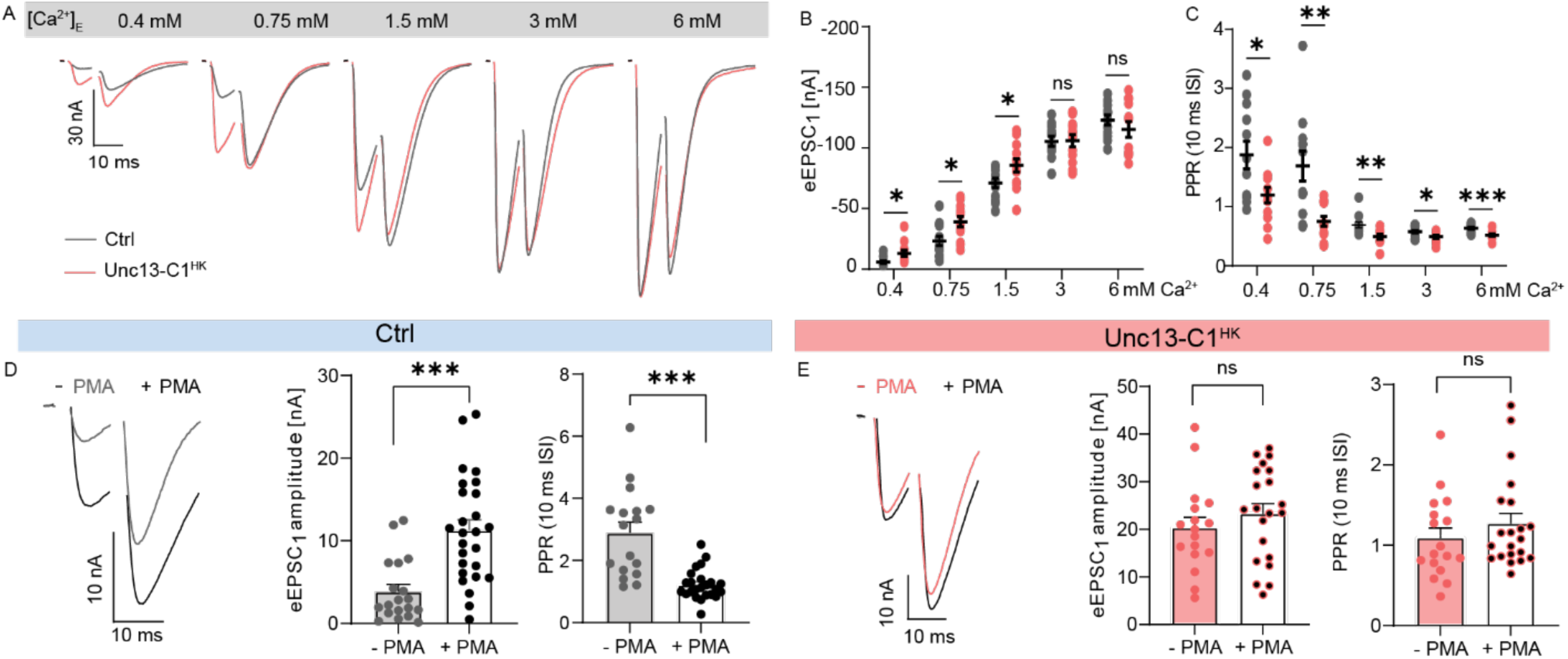
The Unc13-C1^HK^ mutation enhances immediate neurotransmitter release at physiological Ca2+ concentrations and below, abolishes short-term facilitation and blocks phorbol-ester induced potentiation. (**A**) Representative example TEVC traces displaying AP-evoked responses to paired AP stimulation (10 ms interstimulus interval) from a single Ctrl (grey) and Unc13-C1^HK^ (red) NMJ, acquired at progressively increasing extracellular Ca^2+^ concentrations [Ca^2+^]_E_ ranging from 0.4 to 6 mM. Quantification of first (**B**) eEPSC_1_ amplitudes and (**C**) PPRs in Ctrl (grey) and Unc13-C1^HK^ (red) conditions. (**D**) Analysis of control (Ctrl) animals at 0.4 mM extracellular Ca^2+^ concentration. Left: representative traces of AP-evoked paired-pulse responses recorded after a 10-minute preincubation with either DMSO (−PMA) or 2 μM PMA in DMSO (+PMA). Middle: quantification of eEPSC_1_ amplitudes. Right: quantification of Paired-Pulse-Ratios (PPRs, PPR=eEPSC_2_/eEPSC_1_) (10 ms interstimulus interval [ISI]). (**E**) Analysis of Unc13-C1^HK^ animals at 0.4 mM extracellular Ca^2+^ concentration. Left: representative traces of AP-evoked paired-pulse responses recorded after a 10-minute preincubation with either DMSO (−PMA) or 2μM PMA in DMSO (+PMA). Middle: quantification of eEPSC_1_ amplitudes. Right: quantification of PPR ratios (10 ms interstimulus interval [ISI]). Number of cells (n) and animals (N) investigated: n/N: n/N(*Ctrl*, **B** and **C**) = 12/12, n/N(Unc13-C1^HK^, **E**)= 12/12, n/N(*Ctrl* -PMA,**D**) = 19/9, n/N(*Ctrl* +PMA, **D**) = 26/11, n/N(Unc13-C1^HK^ -PMA, **E**) = 17/7, n/N(Unc13-C1^HK^ +PMA, **E**) = 21/7. For exact genotypes see methods. Data depict mean values ± SEM. Statistical analysis with Mann-Whitney U test (B, C, D, E). n.s., p > 0.05; *p % 0.05; **p % 0.01; ***p % 0.001.

A corresponding mutation of the C1 domain in the mammalian ortholog Munc13-1 blocks the potentiation of neurotransmitter release in response to phorbol esters^16^ and we tested if this property was shared at the fly NMJ. Indeed, unlike control cells, where phorbol ester treatment (PMA, 2 μM for 10 minutes) profoundly increased eEPSC_1_ amplitudes and decreased PPRs (Fig. 6D)^66, 67^, no effect was seen in Unc13-C1^HK^ animals (Fig. 6E), indicating a conserved function of this domain. Furthermore, acute application of octopamine enhanced eEPSC_1_ amplitudes measured at 1.5 mM external Ca^2+^ concentrations in wildtype but not Unc13-C1^HK^ mutant animals (Fig. S6.1A and B), demonstrating that octopamine also potentiates neurotransmitter release at physiological Ca^2+^ levels and corroborating a requirement of Unc13 C1 domain for this.

### Organismal consequences of Unc13-C1^HK^ mutation

Considering that the locomotor activity is octopamine dependent^68,69^ we investigated the locomotion behavior of Unc13-C1^HK^ mutants at two different lifetime points. We quantified 3^rd^ instar larvae crawling speed from trajectories of 1-minute duration as well as locomotor activity of adult flies. Unc13-C1^HK^ mutant larvae showed decreased locomotor speeds in comparison to control w1118 larvae (Fig. 7A). Likewise, adult Unc13-C1^HK^ mutants of both sexes displayed reduced locomotor activity throughout the circadian cycle (Fig. 7B, S7C). Previous studies have shown that octopamine is secreted from type II boutons during starvation states which enhanced locomotion speed via the Octβ2R^23^. We wondered whether the pathway described here also contributed to this behavioral adaptation and therefore tested whether this was affected in Unc13-C1^HK^ mutants. Both mutant and wildtype 3^rd^ instar larvae starved for two hours robustly increased locomotor speeds (Fig. 7B), suggesting that this behavioral adaptation does not rely on a regulation of the Unc13-C1 domain. We also tested possible involvement of motoneuronal OAMB receptors by their cell specific knockdown (*OK6-Gal4>UAS-OAMB-RNAi*). OAMB knockdown neither affected crawling speeds in fed animals (Fig. S7.1A) nor did it affect the starvation-induced locomotion (Fig. S7.1C). These results suggest the involvement of a separate pathway, consistent with the known role of Octβ2R/Octβ1R and the cAMP pathway for this starvation induced adaptation^23, 24^.

**Figure 7:**
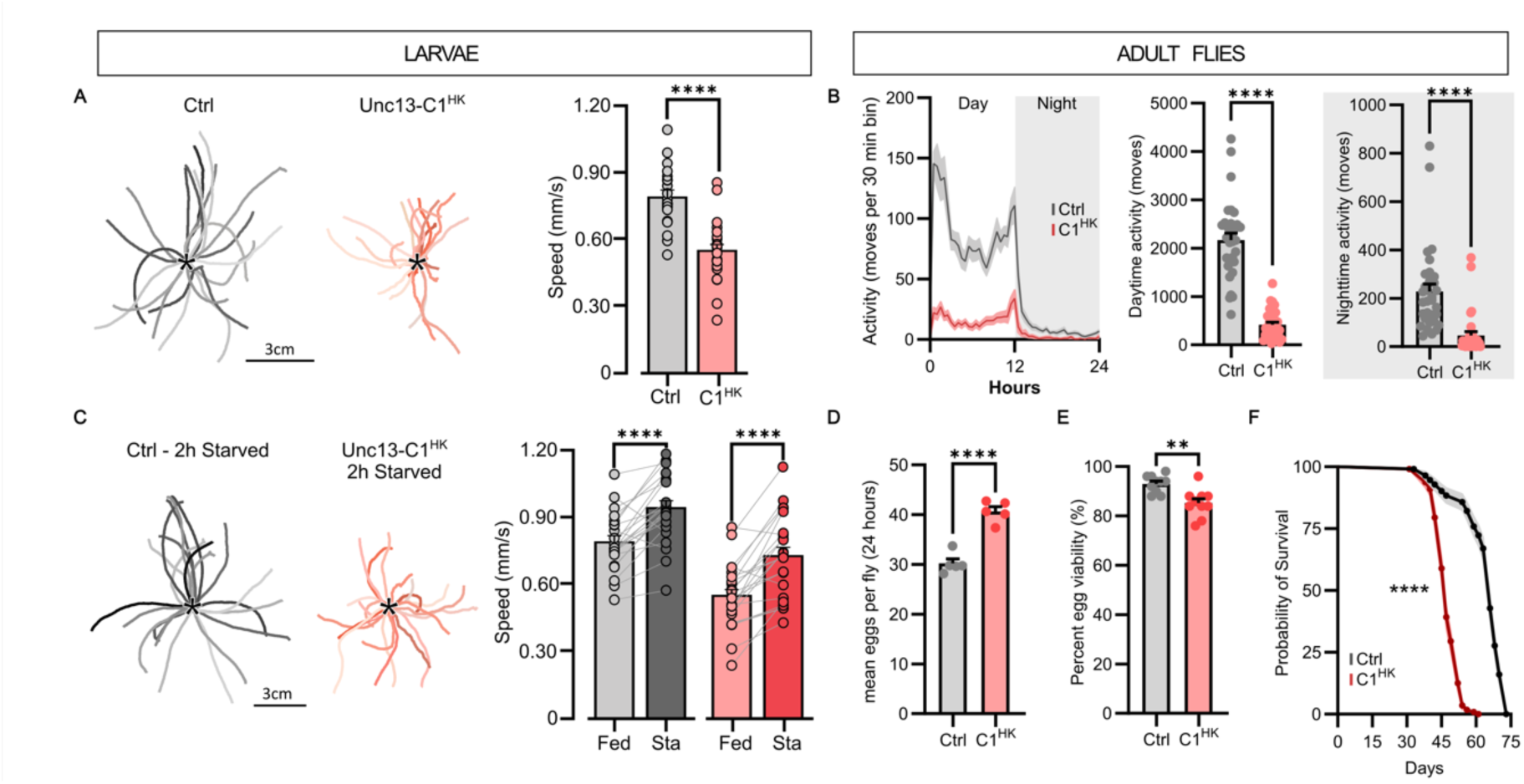
Unc13-C1^HK^ flies show decreased locomotor speed, normal adaptation to starvation, increased reproduction and decreased lifespan. (**A**) 1-minute trajectories of 3^rd^ instar larvae crawling filmed at 15 frames/s (*, starting position) from w1118 larvae (Ctrl, grey traces) and from Unc13-C1^HK^ mutant (C1^HK^, light red traces). In comparison to control larvae (n = 22), crawling speed is diminished in Unc13-C1^HK^ mutants (Number of animals (n): n(Ctrl) = 22, n(C1^HK^) = 22. Unpaired t-test, p = 1e^-06^). (**B**). Locomotor activity profile of adult female control and Unc13-C1^HK^ animals across 24 hours (left) and quantification of total activity split in day (centre) and night (right). Number of animals (n): n(Ctrl) = 32, n(C1^HK^) = 32. Unpaired Mann-Whitney test, daytime activity: p = <0.0001, nighttime activity: p = <0.0001. (**C**) 2h-starvation significantly increases crawling speeds in Unc13-C1^HK^ mutants and in control larvae (C1^HK^: Paired t-test, p = 3.9e^-06^; Ctrl: Paired t-test, p=9.5e^-06^). (**D**) Quantification of fecundity of control and Unc13-C1^HK^ animals. Number of animals (n), number of groups (N): n/N(Ctrl) = 70/5, n/N(C1^HK^) = 70/5. Unpaired t-test, p< = 0.0001. (**E**) Quantification of fertility of control and Unc13-C1^HK^ animals. Number of animals (n), number of groups (N): n/N(Ctrl) = 500/10, n/N(C1^HK^) = 500/10. Unpaired t-test, p< = 0.0014. (**F**) Longevity analysis of female control and Unc13-C1^HK^ animals. Lifespan is reduced in female Unc13-C1^HK^ flies. Number of animals (n): n(Ctrl) = 112, n(C1^HK^) = 112. Log-rank (Mantel-Cox) test, p=<0.0001. Error bars represent ± SEM. **: P <= 0.01; ***: P <= 0.001; ****: p <= 0.0001.

Female reproductive processes including oogenesis, ovulation and adaptive behavior associated with reproduction also critically depend on octopaminergic signaling^70, 71, 72^. We therefore investigated the outcome of the Unc13-C1^HK^ mutation on reproductive output by quantifying fecundity and offspring viability. Young (4-6 days) Unc13-C1^HK^ mutants showed a significantly increased number of deposited eggs (+35 %) over a 24-hour period compared to wildtype controls (Fig. 7D). Moreover, viability of the produced offspring was only slightly reduced in the mutants (−9 %) (Fig. 7E), indicating that overall reproductive output was increased. Finally, since a gain in early-age reproductive output is generally balanced by changes in longevity^73^, we estimated the lifespan of Unc13-C1^HK^ mutants compared to controls. Consistent with a negative trade-off relationship between reproduction and longevity, Unc13-C1^HK^ mutants showed a shortened lifespan (Fig. 7F, S7.1D). Altogether, chronic activation of Unc13 in the Unc13-C1^HK^ mutation compromises survival and locomotion while reproduction is increased. This demonstrates that the ability to modulate Unc13A via its C1 domain is of widespread relevance to the organism.

## Discussion

We here set out to investigate mechanisms of fast neurotransmitter release potentiation. Electrophysiological experiments established that the monoamine octopamine rapidly increases neurotransmitter release at the *Drosophila* 3^rd^ instar larval NMJ within one minute by a presynaptic mechanism. Potentiation depended on PLC activity and while no significant alterations of AP-induced Ca^2+^ influx were seen, the confocal signals of BRP and Unc13 increased with 1-minute octopamine treatment. A dependence on Unc13 for this fast potentiation was corroborated genetically and identified a pivotal role of the DAG binding Unc13 C1 domain. A point mutation in the domain not only blocked the potentiation by octopamine and DAG analogs, but also increased baseline neurotransmitter release, caused short-term synaptic depression, and reduced synaptic Unc13 levels. Live, single molecule microscopy revealed a rapid rearrangement of the Unc13 on the nanoscale coincident with this octopamine induced potentiation via the C1 domain. At the behavioral level, the locomotion and survival of Unc13-C1^HK^ mutants flies were considerably reduced, while the reproduction was enhanced.

In our NMJ analysis we exclusively focused type I larval NMJs at muscles 4 and 6. However, these muscles are not contacted by the peptidergic type II NMJs known to release octopamine at other muscles (muscle 12)^23, 47, 74^. We attempted visualization of local octopamine reservoirs at type I NMJs by immunostaining, but failed to see clear structures, possibly due to low levels, unspecific background staining or a widespread distribution of octopamine throughout the tissue. The *Drosophila* genome encodes five different receptors, three beta-adrenergic-like Receptors Octβ1R, Octβ2R and Octβ3R as well as the two alpha-adrenergic-like receptors OAMB and Octα2R which have widespread distribution throughout the nervous system^75^. At larval NMJs, acute octopamine application was seen to elevate cAMP levels (of muscle 12), trigger neuropeptide release (from muscle 6/7), and enhance glutamate release, at least partly by Octβ1R and Octβ2R dependent cAMP signaling^23, 42^. Recently, the expression levels of the OAMB receptor were assessed at type I NMJs by RNA seq which reported similar levels as for Octβ1R and Octβ2R, and similar expression in the small and big type Is and Ib boutons^50^. We here show that presynaptic OAMB receptor knockdown fully blocked potentiation of neurotransmitter release by 1-minute octopamine treatment in current clamp recordings (Fig. 1B), indicating that this receptor is needed for fast potentiation. OAMB has two isoforms, OAMB-K3 and OAMB-AS, which differ in signaling pathways: OAMB-K3 increases both Ca²⁺ and cAMP, while OAMB-AS only elevates Ca²⁺, implicating the involvement of the PLC pathway and in some cases activation of CaMKII^76, 77, 78^. Both isoforms (OAMB-K3 and OAMB-AS) regulate various physiological processes, including learning, consistent with a relevance of Gαq/PLC signaling alongside the established Gαs/Gαi/cAMP pathway in these processes.

We saw that although fast octopamine-induced potentiation was fully blocked by pharmacological PLC inhibition (Fig. 1C), its knockdown in presynaptic motor neurons merely attenuated it (Fig. S1.2G and H). This residual effect could be due to incomplete gene silencing or the involvement of other cell types. In further support of the PLC pathway, mutation of the DAG-binding Unc13 C1 domain fully blocked potentiation (Fig. 4I, J and S6.1B) pointing to a direct role of DAG signaling lipid in this fast neuromodulation. We cannot exclude additional downstream targets for PLC contributing to synaptic potentiation and indeed expect that these will be needed on the longer term. For instance, PLC activity also generates intracellular IP_3_ which releases Ca^2+^ from intracellular stores and has a function in long-term homeostatic potentiation^79^. One possibility is that this involves activation of the Unc13 C2B domain, whose mutation also partially blocked octopamine-induced potentiation (Fig. 4G and H). Additionally, apart from directly activating the Unc13 C1 domain, DAG activates protein kinase C (PKC), which has several phosphorylation targets in the neurotransmitter release machinery, including (M)Unc18, Synaptotagmin-1 and SNAP25 whose phosphorylation affect neurotransmitter release^32, 33, 80^. Future work should focus on establishing which signaling pathways and messengers mediate distinct temporal phases of synaptic plasticity.

PMA strongly and rapidly potentiates neurotransmitter release via the (M)Unc13 C1 domain^16, 17, 66, 67^. Our results align with this since the same amino acid exchange in the fly ortholog also blocked PMA induced potentiation at the NMJ (Fig. 6). We here provide single molecule analysis of this Unc13-C1^HK^ mutant and a comparison between mutant and wildtype synapses revealed a severe reduction in the number of Unc13 molecules assessed by bleach curve analysis (Fig. S2.3 and S5.1). Notably, lower confocal signals of antibody-labelled Unc13 were observed for mutants of the Ca^2+^ calmodulin binding domain, the C2B and the C1 domain which all had increased baseline responses and lower capacity to be potentiated by PMA and/or octopamine (Fig. 4E and I; Fig. 6E; ^66^). While it is typically observed that AZ protein levels increase with potentiation, this indicates the opposite: Lower protein levels in cases where baseline transmission is enhanced, opening the interesting possibility of bi-directional homeostasis. Consistent with this, the overall effects of the mutations were low at physiological external Ca^2+^ concentrations (Fig. 6)^66^, which could illustrate an attempt to normalize synaptic communication in this range.

Besides disrupting octopamine-induced potentiation, the Unc13-C1^HK^ mutation also caused short term synaptic depression. Moreover, our behavioural analysis points to consequences at the level of the full organism. A marked decrease in locomotor speed was seen both in larval and adult flies (Fig. 7). This is consistent with reported locomotor defects in *C. elegans* and even human patients with Unc13 point mutations^81, 82^. Loss of octopaminergic input also decreases larval crawling speeds, but this is unlikely to rely on OAMB-dependent signalling at the level of type I motoneurons since we observed no effect of its knockdown (Fig. 7.1). Given the marked change in short-term plasticity, we predict that this mutation likely affects the temporal pattern of motoneural activation, which is consistent with the observed disruption of coordinated movement in *C. elegans* and human patients^81, 82^. Future research investigating the effect of this mutation on the central pattern generator for locomotion may thus provide valuable insight into the relevance of Unc13 modulation for coordinated movements.

Previously reported changes of the synapse structure typically unfold over much longer timescales, for instance, prolonged changes by octopamine described in^24^, where synaptic growth occurs over several days. On a comparable timescale, long term potentiation accumulated Munc13 and shortened coupling distances to presynaptic Ca^2+^ channels in the mammalian hippocampus^83, 84^. Changes in alignment between presynaptic Munc13 and postsynaptic receptors nanoclusters were seen on a timescale of 25 minutes^85^ and on a similar timescale, live single molecule imaging of calcium channels and BRP at the fly larval NMJ described their nanoscopic compaction during sustain phase of PHP^11^. Several studies described structural changes during PHP on a faster timescale of 10-minutes^7, 8, 58^. Here we found increased Unc13A confocal signals (Fig. 1E) following only 1-minute octopamine treatment. This result was corroborated by live single molecule sptPALM which demonstrated an instant decrease in Unc13 motility, followed by compaction of Unc13 signal distribution within the AZ (Fig. 2). During this 1-minute octopamine-induced structural reorganization, no changes in the relative number of Unc13 molecules were observed (Fig S2.3A-C) while undergoing this structural redistribution. Perhaps, this reflects the first stage of acute potentiation that reorders and compacts AZ components, as previously described after 10-minute PHP^86^ which then is further consolidated by an increases in relative numbers of AZ proteins, as is observed for BRP and Cac scaffold proteins after 20-minutes of PHP^11^. Meanwhile, the decrease in dynamics observed in Figure 2 could be caused by the crowding of molecules at the AZ, especially as related proteins like BRP and Ca^2+^ channels might simultaneously undergo similar enrichment^11^.

A picture is emerging where slowed diffusion and nanoscopic compaction of multiple proteins coincides with neurotransmitter release potentiation. How exactly this enhances synaptic output is not clear yet. One explanation could be that simultaneous “freezing” of Ca^2+^ channels and scaffolding proteins with vesicular release sites could lock those reaction partners in space to maintain a robust coupling distance for Ca^2+^-induced release^87, 88^. A direct effect of the here observed local Unc13 concentration on the nanoscale could relate to its essential role in tethering SVs to the plasma membrane^89^. We estimate from our bleach curve analysis that merely 8-17 Unc13 molecules are present per AZ on average (Fig. S2.3). Transmitter release is limited to few release sites per AZ and assuming an average of 4 sites per AZ, this corresponds to only ∼3 Unc13s per site at rest^7, 55, 90, 91^. This relatively small number already indicates that successful SV capture will be rare, especially if multiple Unc13s need to cooperate in successful reactions. The other major limiting factor is space: Due to the small SV size, the nanoscopic distribution of Unc13 greatly matters, for instance to ensure that multiple proteins would be within an SV’s reach. Accordingly, we here demonstrate that both critical parameters adapt with potentiation: The seen compaction indicates more Unc13 proteins occupy a smaller area (Fig. 2N & P). (M)Unc13 was suggested to energetically stabilize SV priming^92^ and we here attribute the observed potentiation to an enlargement of the readily releasable pool of vesicles consistent with this. Whether this causes morphological changes that can be detected in the synaptic ultrastructure remains to be established. On the one hand, previous analysis of distinct mammalian central synapses with low/high release output and high/low phorbol ester enhancement attributed this to distinct primed states without detectable differences in SV docking^93^. On the other hand, recent cryo-electron tomography of mammalian synaptosomes treated with phorbol esters reported small alterations in SV docking distributions and increased SV tethering, arguing that differences might be resolvable under certain conditions^89^. More work will be needed here to figure out whether these different physiological states can indeed be distinguished morphologically.

We used the *Drosophila melanogaster* NMJ as a model system owing to its unmatched accessibility for live single molecule imaging, genetic manipulation, electrophysiological recordings and behavioral profiling. However, due to the fundamental role of the PLC pathway we predict that the fast potentiation mechanism described here holds relevance throughout the nervous system and across species. Indeed, α1-adrenergic receptors (α1-ARs), the corresponding receptors of OAMBs in mammals, mediate short-term plasticity, long term potentiation and influence cognition, learning and memory^94, 95, 96^. α1A-AR receptors were suggested as potential drug therapeutic for conditions the treatment of neurodegenerative diseases like Alzheimer’s disease^97^. Future work based on the insight provided here could test the generality of this mechanism and its intersection with human disease.

## Material and Methods

### Fly Husbandry, Stocks, and Handling

Please see Key Resource Table (Table 1) above for full details of these strains. Fly strains were maintained under standard laboratory conditions ^98^ and raised at 25 degrees Celsius. For experiments, male and female third instar larvae (larval stage 3) were used unless otherwise stated. The genotypes utilized included Wild-type: +/+ (w1118), sourced from the Bloomington Drosophila Stock Center/WellGenetics; OK6-Gal4 ^99^; UAS-Unc13A-RNAi ^56^; UAS-OAMB-RNAi ^100^; UAS-syt-mScarlet-GCaMP8f ^101^; UAS-PLC-RNAi ^102^; Unc13 Biosensor ^56^; *Unc13A C2B^K1862W^* (this lab); *Unc13A C1^H1723K^* (this lab); *Unc13mOES3.2* (this lab); *Unc13C1^H1723K^ mOES3.2-*(this lab)

**Table 1:**
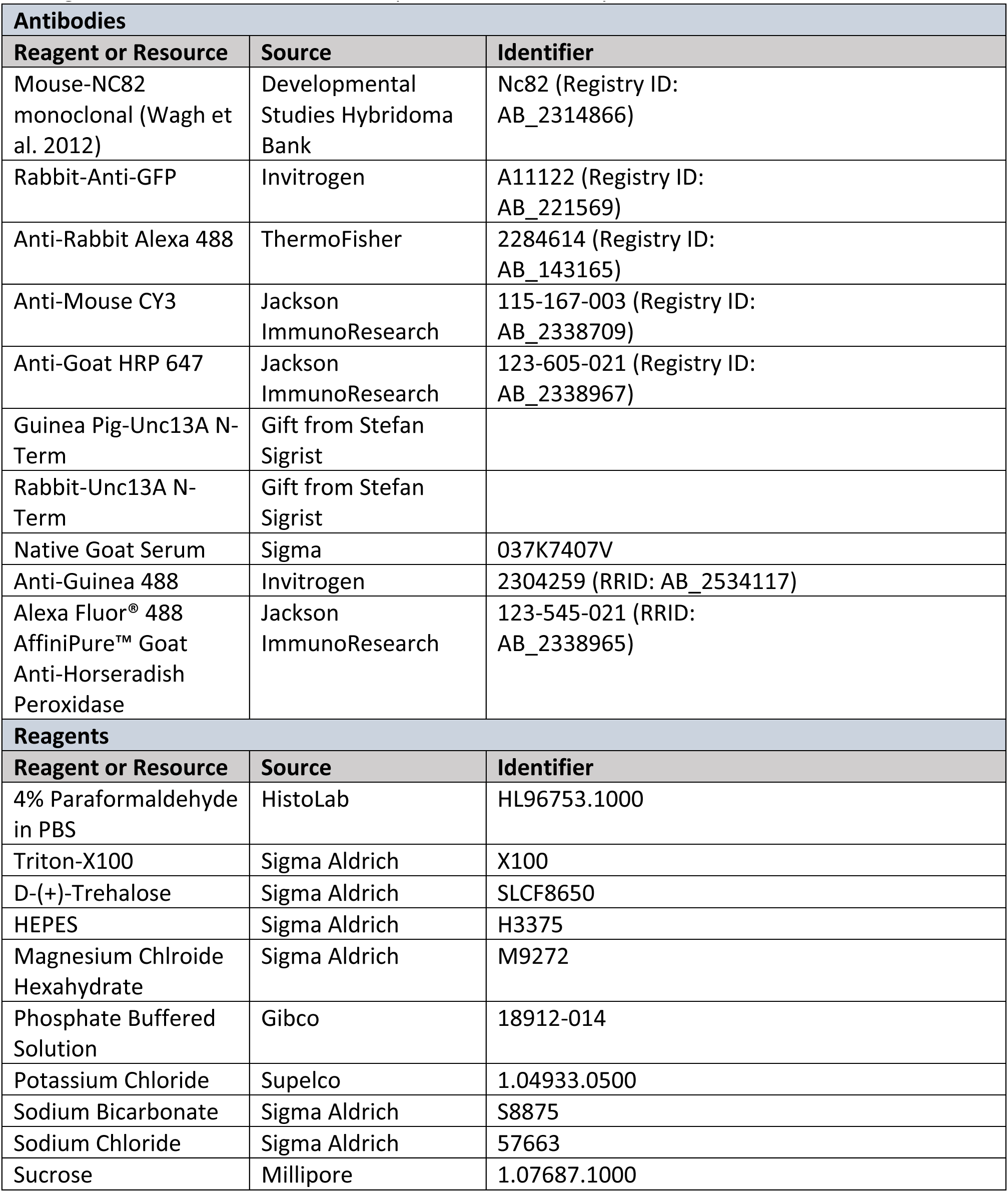
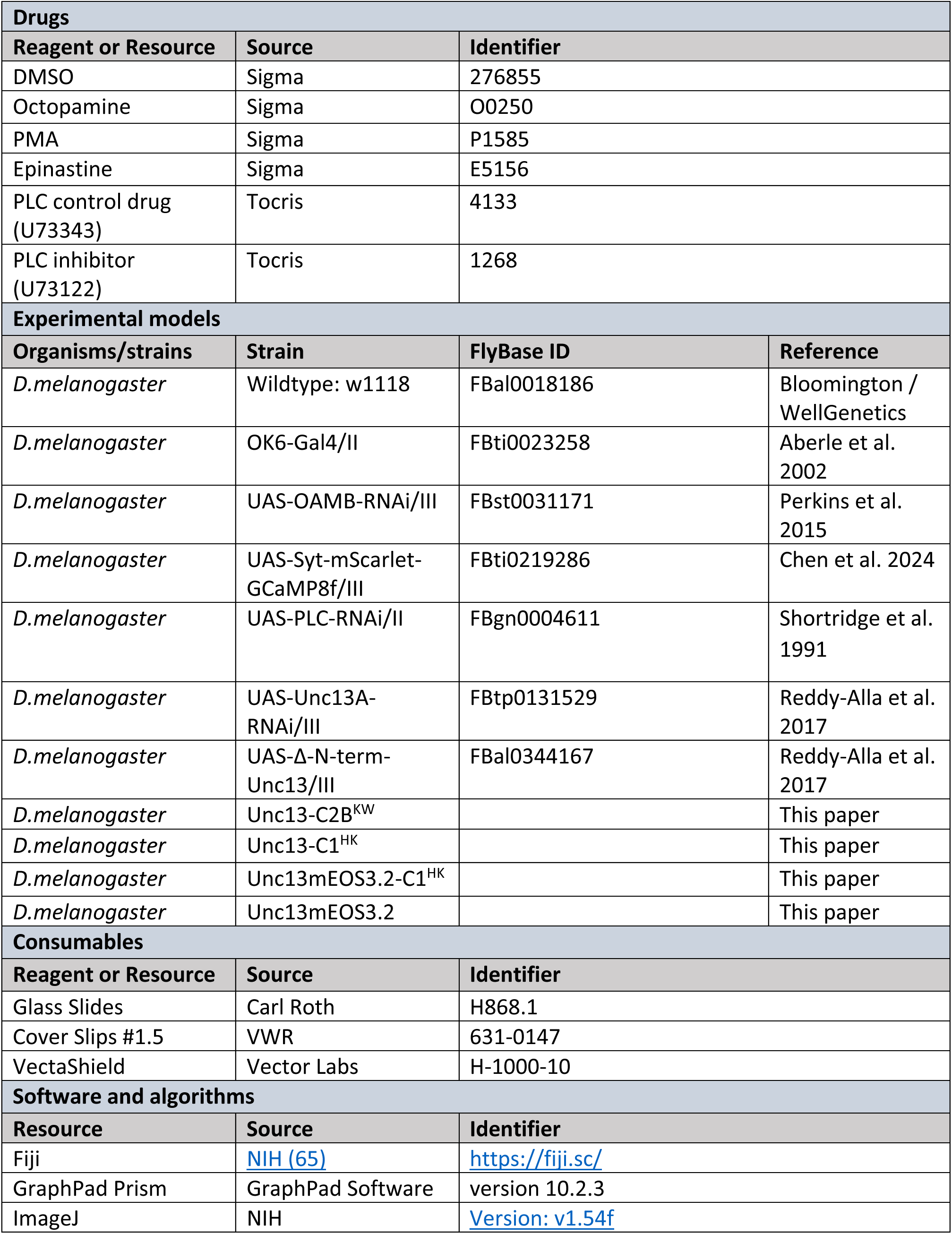
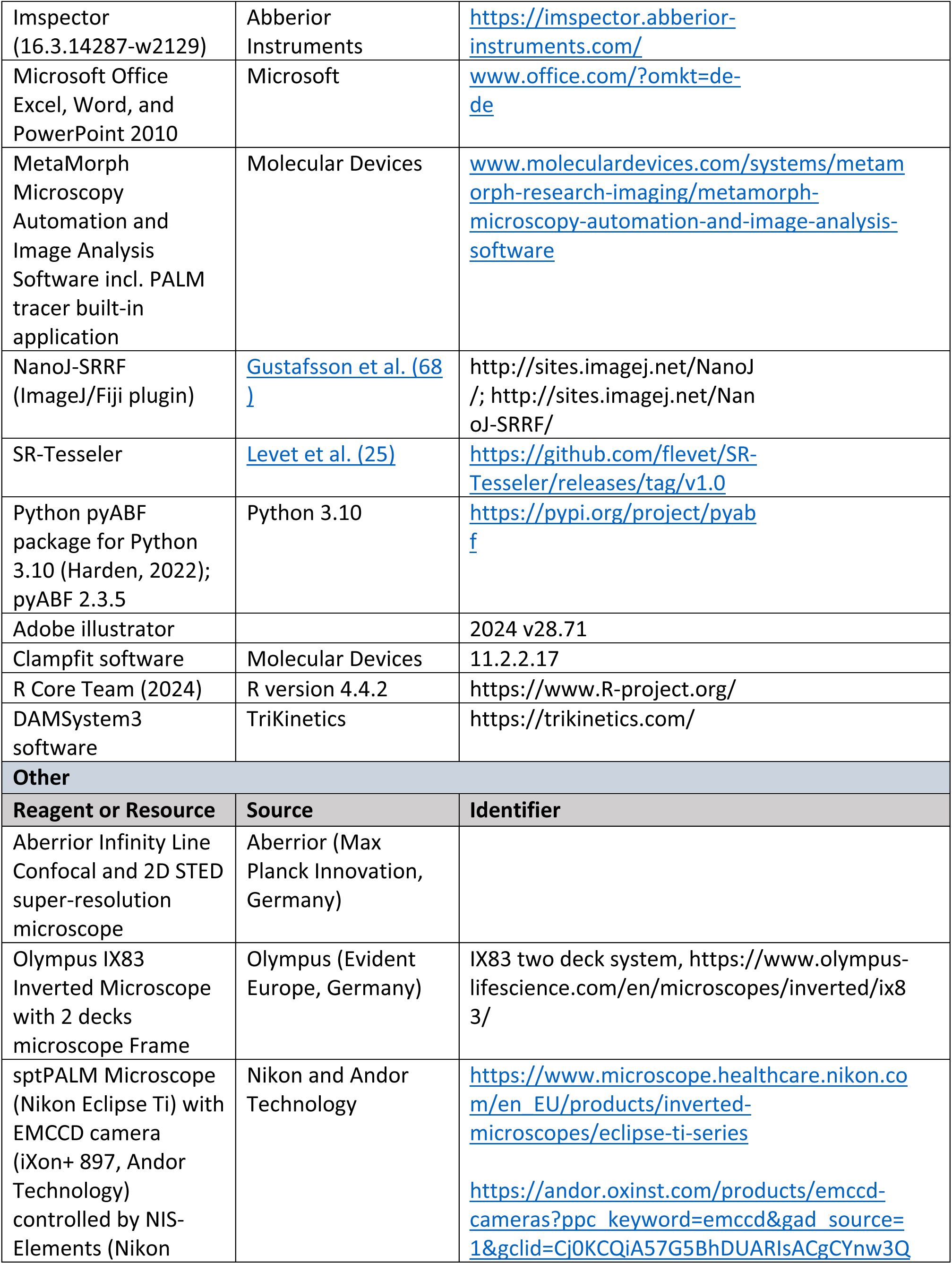
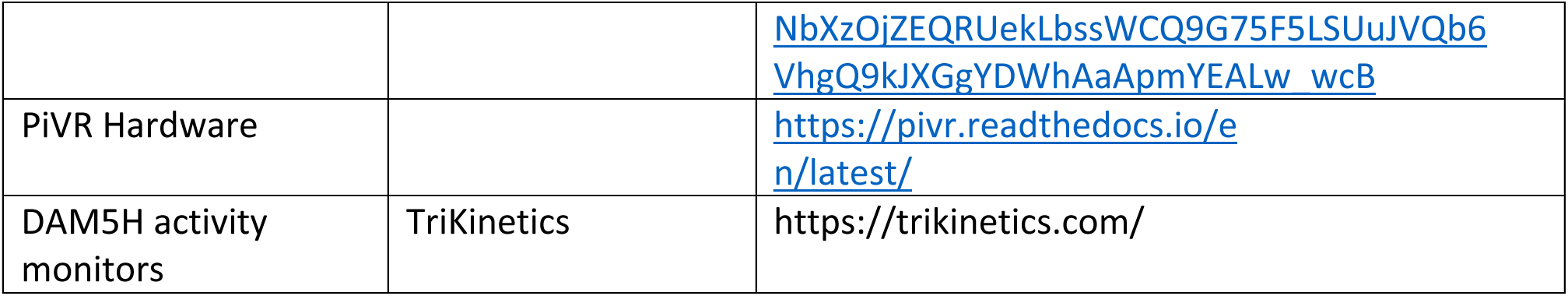
Key resources. Table provides an overview of the resources employed in this study, including antibodies, reagents, drugs, *Drosophila melanogaster* strains, consumables, software and algorithms and other resources important for this study.

#### Genotypes

**Fig. 1**: Ctrl(A & E): w1118, OAMB KD(B): *;OK6-Gal4/+;UAS-OAMB-RNAi/+;,* PLC inhibitor(C): *w1118,* Syt::GCaMP8f (D): ; *OK6-Gal4/+;UAS-syt-mScarlet-GCaMP8f/+;*.

**Fig. S1.1**: Ctrl (A): *w1118*, OAMB ctrl(F,H): ;;*UAS-OAMB-RNAi/+;,* OAMB KD(G,H)*: ;OK6-Gal4/+;UAS-OAMB-RNAi/+;,* naïve controls(E): *;OK6-Gal4/+;UAS-syt-mScarlet-GCaMP8f/+*.

**Fig. S1.2**: Ctrl Drug/PLC Blocker (B-D, I-K): *w1118*, Ctrl(F-G,L-N)*: ;UAS-PLC-RNAi/+;;*, PLC knockdown (F-G, L-N): *;UAS-PLC-RNAi/OK6-Gal4;;*.

**Fig. 2**: Unc13mEOS3.2

**Fig. S2.1**: *w1118*, *Unc13mEOS3.2*

**Fig. S2.2**: *Unc13mEOS3.2*

**Fig. S2.3**: *Unc13mEOS3.2*

**Fig. 3**. Genotypes: Ctrl(B-D): ;;*UAS-Unc13A-RNAi/+;*, Unc13A KD (B-D): *;OK6-Gal4/+; UAS- Unc13A-RNAi/+;,* Unc13 Biosensor(F,G): *;OK6-Gal4/OK6-Gal4;UAS-Δ-N-term-Unc13/UAS-Δ-N-term-Unc13;*

**Fig. S3.1**: genetic ctrl(A,C): ;;*UAS-Unc13A-RNAi/+;,* Unc13A KD(B,C): *;OK6-Gal4/+; UAS-Unc13A- RNAi/+;*.

**Fig. 4**: Ctrl(B-J): *w1118,* C2B^KW^(B,C,G,H): ;;;*Unc13-C2B^K1862W^*/ *Unc13-C2B^K1862W,^* C1^HK^(E,F,I,J): ;;;*Unc13-C^H1723K^/ Unc13-C^H1723K^*.

**Fig S4.1**: Ctrl(A-L): *w1118,* C2B(A-D,I,J): *Unc13-C2B^K1862W^*, C1(E-H,K,L): *Unc13-C1^H1723K^*.

**Fig. 5**: Unc13mEOS3.2: *Unc13mEOS3.2,* Unc13-C1^HK^mEOS3.2: *Unc13-C1^H1723K^mEOS3.2*

**Fig. S5.1**: Unc13mEOS3.2: *Unc13mEOS3.2,* Unc13-C1^HK^mEOS3.2: *Unc13-C1^H1723K^mEOS3.2*

**Fig. 6**: Ctrl: *w1118,* Unc13-C1^HK^: *Unc13-C1^H1723K^*

**Fig. S6.1**: Ctrl: *w1118,* Unc13-C1^HK^: *Unc13-C1^H1723K^*

**Fig. 7**: Ctrl: *w1118,* Unc13-C1^HK^: *Unc13-C1^H1723K^*

**Fig. S7.1**: OK6>w1118: *;OK6-Gal4/+;;,* OK6>OAMB-RNAi: *;OK6-Gal4/UAS-OAMB-RNAi;;*, Ctrl: *w1118*, C1^HK^: *Unc13-C1^H1723K^*

### Fly mutagenesis Unc13 C1^HK^ and Unc13-C2B^KW^

The Unc13 c1 domain mutation was developed based on the analogous change made in mouse Munc-13-1 homologue, where histidine 567 was replaced by lysine. By aligning sequences, histidine 1723 was identified as the corresponding target residue in *Drosophila* Unc13A. Since both Unc13A and Unc13B splice variants possess a C1 domain ^57^ this single alteration affects both variants. CRISPR-mediated mutagenesis was conducted by WellGenetics Inc. following modified methods from Kondo and Ueda ^103^ to introduce a targeted break in unc-13/CG2999. For repair, A PBacDsRed cassette containing two PBac terminals and 3xP3-DsRed and two homology arms with point mutations (one inducing the amino acid change and one introducing a TTAA sequence to recognize proper integration on the other homology arm) was employed. Initial screening of F1 progeny used the DsRed marker to identify successful integration events. Afterwards, the marker was excised and PCR plus sequencing of genomic DNA confirmed the presence of the desired point mutation and the complete removal of the cassette in the final *Drosophila* line ^104^.

The flies used for experiments were validated using genomic PCR and sequencing methods to verify PiggyBac transposition alleles of *w[*];;; unc13-PA H1723K CRISPR[PBacDsRed] / In(4)ciD, ciD panciD* fly by testing if the selection marker(3XP3-DsRed) is precisely excised, the HK mutation present and an integrated TTAA sequence left behind in the coding exon of *unc- 13/CG2999.* For the Unc13-C1^HK^ mutant the forward primer 5’-CCGCTACAAAACGAAATGCT targeting the upstream homology arm the reverse primer, 5’-GTCCATAACAAAAAATTCTT, targeting the downstream homology arm were used to verify the mutation in flies used for experiments.

### Generation of on-locus tagged Unc13-mOES3.2 and Unc13-C1^HK^-mEOS3.2

Briefly, C-terminally tagged *Unc13mOES3.2* was generated using a scarless CRISPR/piggyBac-based approach (https://flycrispr.org/). Tag mEOS3.2 and an assigned linker were knocked in right before the Stop Codon of Unc13, at the same stable integration site as the previously published Unc13-GFP construct ^56^.

The transgenic flies were produced by Well Genetics Inc. (Taipei City, Taiwan), following adapted protocols from Kondo and Ueda ^103^. Specifically, these lines were generated through CRISPR-Cas9–mediated genome editing via homology-dependent repair, employing a single guide RNA (gRNA) and a double-stranded DNA plasmid donor.

CRISPR endogenous tagging of *w[*];;; unc13-PA H1723K CRISPR[PBacDsRed] / In(4)ciD, ciD panciD* at C-terminal insertion position before the stop codon (position shared in Unc13mEOS3.2) is tagged with mEOS3.2 to produce of *w[*];;; unc13mEOS3.2-PA H1723K CRISPR[PBacDsRed] / In(4)ciD, ciD panciD and* testing if the selection marker(3XP3-DsRed) is precisely excised, the HK mutation present and an integrated TTAA sequence left behind in the coding exon of *unc-13/CG2999.* For the Unc13mEOS3.2-C1^HK^ mutant the forward primer 5’-CCGCTACAAAACGAAATGCT targeting the upstream homology arm the reverse primer, 5’-GTCCATAACAAAAAATTCTT, targeting the downstream homology arm were used to verify the mutation in flies used for experiments.

### Tissue preparation and immunohistochemistry for confocal microscopy

3^rd^ instar larvae of interest were first pinned at the head and tail onto a dissecting pad and covered with Ca^2+^ free and low Mg^2+^ Hemolymph-Like Saline Solution 3 (HL3). They were then dissected dorsally along the midline, the dorsal walls were lightly stretched and pinned to the pad, and the organs and central nervous system were removed. Larvae would undergo drug treatments or, if no drug treatments, immediately fixed in 40 µL of 4% paraformaldehyde in phosphate buffer saline (PFA, HistoLab, Askim, SE, HL96753.1000) at room temperature for 10 minutes. Larvae would be washed with 30 µL phosphate buffered saline solution with 0.05% Triton-X100 (PBST, Sigma Aldrich, MO, USA, X100). After washing every 20 minutes for 3-5 times in PBST, larvae would be blocked for 1 hour at room temperature in normal goat serum (NGS, 5% solution volume, Sigma Aldrich, MO, USA, 037K7407V) diluted in PBST. Larvae would then be incubated overnight or over the weekend with primary antibodies: rabbit-Unc13A N-term (1:250, Fig. 4C and 4F, ^57^) or rabbit anti-GFP (1:250, Fig 3F and 3G, Thermo Fisher Scientific, MA, USA, A11122; RRID: AB_221569) or guinea pig-Unc13A N-term (1:250, Fig. 1E^57^), and mouse-nc82 (1:250, Developmental Studies Hybrodoma Bank [HSDB], University of Iowa, Iowa city, IA, USA; RRID: AB_2314866). The next day, larvae were washed 3 times consecutively in PBST, and then once every 20 minutes in PBST 3-5 times. Larvae were then incubated in PBST, NGS (5% solution volume) and secondary antibodies in the following concentrations: goat anti-rabbit Alexa Fluor 488 (1:500, Fig. 4C and 4F, Thermo Fisher Scientific, MA, USA, A11008; AB Registry ID: AB_143165), or Alexa Fluor 488 goat anti-guinea pig (1:500, Fig. 1E, Thermo Fisher Scientific A11073; RRID: AB_2534117), goat anti-mouse CY3 (1:500, Fig. 1E, 4C and 4F, Jackson ImmunoResearch, PA, USA, 115-167-003; AB Registry ID: AB_2338709), goat anti-HRP Alexa Fluor 647 (1:250, used in imaging experiments in Fig. 1,3,4, not shown), Jackson ImmunoResearch, PA, USA, 123-605-021; AB Registry ID: AB_2338967). After incubation, larvae were washed 3-5x in PBST every 20 minutes. For confocal imaging, larvae were mounted in VectaShield (Vector labs, CA, USA H-1000-10) on a microscope slide and sealed with nail polish under a cover slip (Carl Roth, Germany, H868.1).

#### Octopamine treatments for confocal analysis

3^rd^ instar larvae were incubated with 30 µL of either 20 µM octopamine for 1 minute, 2 µM PMA for 10 minutes, or DMSO (0.1%) for 1 minute.

#### Epinastine treatments for confocal analysis

3^rd^ instar larvae were pre-incubated for 5 minutes in 30 µL of either 0.1% DMSO for both octopamine and DMSO groups, or 1 µM epinastine. These larvae would then be further incubated by either 0.1% DMSO or 20 µM octopamine (final concentration) for 1 minute. This final octopamine concentration was achieved by making a solution with 40 µM octopamine diluted in HL3 and another solution with 40 µM octopamine and 1 µM epinastine in HL3. Therefore, when 30 µL of solution was pipetted onto the larvae, the concentration of octopamine would be halved.

#### Image acquisition, processing, and analysis – confocal microscopy

First the confocal microscope, an Abberior Infinity Line confocal and 3D Stimulation Emission Depletion (STED) super-resolution microscope, was turned on one hour prior to imaging to allow the machinery to warm up. A Plan-achromat oil immersion objective with 63x magnification and a numerical aperture of 1.4 and a 0.19 mm working distance was used. The temperature of the room was maintained around 22 degrees Celsius. A drop of oil was placed onto the cover slip. A larva in the control group was imaged first to preset excitation laser power. Segments A3-A5 muscle 4 NMJs were visualized in live mode to adjust excitation laser channels for Alexa647, CY3, and Alexa488 were adjusted to receive the best resolution without losing the staining due to bleaching. A stack of images was collected at different z positions 300 nm apart in order to capture the entirety of the NMJ, the resolution would be set to 90 nm pixel size with the accumulation set to 3. A total of up to four different NMJs were collected per animal.

#### Analysis of active zone morphology from confocal images at the 3^rd^ instar NMJ

From each z stack, a single image was generated for each investigated channel by creating a projection of the maximal intensities using the Z project function in Image J (version 2.14.0/1.54f). To restrict the analysis of fluorescence signals within the NMJ, the HRP channel was first used to encompass the outline of the NMJ and to create a binary NMJ mask which was then applied to the images obtained in the BRP and Unc13A channel. BRP was used to identify AZs and to create a “region-of-interest” (ROI) mask utilizing a segmentation analysis function, around each AZ at the NMJ^105^. To ensure that each AZ was segmented, each AZ was observed to ensure segmentation function found each AZ. The fluorescence levels in the BRP channel were assessed by finding the mean gray intensity within each ROI. The BRP ROIs were then used as a mask, and then placed on the max projected Unc13A image, and the ROIs were used to assess mean gray intensities of Unc13A in the ROIs. Intensity values were averaged across all AZs in a single NMJ to obtain a representative measure per NMJ. These values were then compared across genotype and treatment conditions.

#### Analysis of the C-terminal Unc13-GFP fragment via line profile analysis from confocal images

Analysis was performed on single confocal imaging slices within the z stack. The specific slice chosen for analysis was made by placing a linear ROI over a single bouton and observing the plot profile of ImageJ in live mode to evaluate the pixel intensity in the GFP channel. The z position within the image stack was moved until the highest pixel intensity in for this ROI was found in the stack. The gray value from the edges of the boutons would be averaged, as these represent Unc13 proteins translocated at the plasma membrane, and therefore the “max” value, as these were observed as the highest intensities in the PMA and octopamine conditions (Fig. 3F, G; filled arrows). The lowest gray value within the cytosol of the bouton would be documented, and therefore the “min” value, as the cytosol of the bouton had generally lower intensity values observed in PMA and octopamine groups (Fig. 3F, G; hollow arrow). Then the bouton’s plasma membrane averaged fluorescence intensity value would be divided by the lowest fluorescent intensity within the same bouton in the cytosol, and this would create a Max/Min ratio. As stated, this ratio could be observed as PMA and octopamine conditions, there was generally a lower fluorescence within the bouton, which enabled us to generate a quantitative measure to compare the degree of translocation of the Unc13 fragment towards the plasma membrane. The rest of the boutons of the NMJ would be taken in a similar manner to get the best representation of the NMJ. Each bouton would be assessed once, and all the boutons would be assessed within an NMJ, only if there were four or more boutons to choose from. Bouton values would then be averaged across their respective NMJ which would be shown as individual points in the results. Each larva would have three NMJs assessed, to give the best representation of the larvae. The data points represented are 1 NMJ, and therefore each animal generally has 3 NMJs on the data set.

### Electrophysiology: Setup and data acquisition

#### Current clamp recordings

All current clamp experiments were performed using third-instar larvae dissected on Sylgard (184, Dow Corning, Midland, MI, USA). Preparations were made in Ca^2+^-free low Mg2+ HL3 (pH adjusted to 7.2 ^7^: 70 mM NaCl, 5 mM KCl, 10 mM MgCl2, 10 mM NaHCO3, 5 mM Trehalose, 115 mM D-Saccharose, 5 mM HEPES) and the dissection procedures followed established protocols^106, 107^. Dissected larvae were then relocated to a recording chamber containing HL3 solution with 0.4 mM Ca^2+^.

Electrophysiological current clamp measurements were obtained at 22.5°C from muscle 6 of abdominal segments A2 and A3. Sharp borosilicate glass electrodes (0.86 x 1.5 x 80 nm, with filament, from Science Products, Hofheim, Germany) were pulled with a P97 pipette puller (Sutter Instrument, CA, USA). The electrodes, backfilled with 3 mM KCl solution, exhibited resistances between 20 and 40 MΩ. Signals were recorded with a Digidata 1440A digitizer (Molecular Devices, Sunnyvale, CA, USA) and Clampex (v10.6) software, and an Axoclamp 900A amplifier (Axon Instruments, Union City, CA, USA) and Axoclamp software. Acquisition was performed at a sampling frequency of 20 kHz with a 5 kHz low-pass filter. Cells with resting membrane potentials higher than −49 mV and input resistances below 4 MΩ before measurements were excluded from the datasets. The data were processed using Clampfit software (v11.2.2.17 ^66^). Graphs present mean values ± S.E.M and were generated using GraphPad Prism 10.1.1 and Adobe Illustrator (Adobe Systems, San Jose, CA, USA).

For data analysis, the mEPSP traces were post-processed with a 500 Hz Gaussian lowpass software filter. A control cell served as the reference for generating a mEPSP template, which was then used to identify events over the entire 30-second recording period. For mEPSP frequency [Hz], the total number of events was then divided by 30 seconds. To assess changes in mEPSP frequency post-treatment, the fold change was calculated by dividing the post-treatment mEPSP frequency by the baseline mEPSP frequency. The eEPSP amplitudes were determined by averaging all 5 stimulated responses in each cell. The fold change in eEPSP amplitudes was calculated by dividing the post-octopamine treatment eEPSPs by the baseline eEPSPs, enabling quantification of the relative change in eEPSPs induced by octopamine treatment.

#### Two Electrode Voltage Clamp

Electrophysiological TEVC experiments were conducted at RT from muscle 6 of abdominal segments A2 and A3. Sharp borosilicate glass electrodes with filaments (0.86 × 1.5 × 80 nm, Science Products, Hofheim, Germany) were prepared using a P97 pipette puller (Sutter Instrument, CA, USA), filled with 3 mM KCl, and exhibited resistances of 20–40 MΩ. Signal acquisition employed a 5 kHz lowpass filter at a sampling frequency of 20 kHz using the Digidata 1440A digitizer (Molecular devices, Sunnyvale, CA, USA) with Clampex (v10.6) software and an Axoclamp 900A amplifier (Axon instruments, Union City, CA, USA) with Axoclamp software ^66^. Cells with resting membrane potentials greater than −49 mV and resistances below 4 MΩ prior to measurements were excluded from further analysis. During all TEVC recordings, the postsynaptic muscle cell was held at −70 mV and only cells maintaining absolute leak currents below 10 nA throughout individual experiments were included into the analysis. The collected data was analyzed using Clampfit software (11.2.2.17).

#### Drug treatments - electrophysiology

For all current clamp experiments, miniature excitatory postsynaptic potentials (mEPSPs) were recorded for 30 s prior to provoking evoked EPSPs (eEPSPs) in current clamp. eEPSP responses were elicited by 5 stimulation pulses (300 µs, 8 V, 0.1 Hz) applied to the respective nerve.

#### DMSO control treatment - electrophysiology (Fig. S1)

After the initial current clamp recording the same larvae were treated for 1 minute with 0.1% DMSO (stored at –20°C).

#### Octopamine treatment - electrophysiology (Fig. 1, S1.1, S1. 2, S2.1, 3, S3.1, 4, S4.1)

After the initial recordings (or following other drug treatments), the same larvae were treated for 1 minute with 20 µM octopamine (stored at –20°C as a 20 mM stock in DMSO). mEPSPs were recorded for 30 seconds prior to evoking eEPSPs in current clamp.

#### PLC Inhibition - electrophysiology (Fig. 1, S1.2)

Larvae were treated for 1 minute with 1 µM PLC inhibitor (Tocris Bioscience, U73122; freshly prepared as a 1 mM stock in DMSO) or with 1 µM PLC control drug (Tocris Bioscience, U73343; freshly prepared as a 1 mM stock in DMSO). Subsequently, mEPSPs and eEPSPs were recorded following octopamine treatment in current clamp.

#### TEVC recordings of baseline transmission and short-term plasticity in Unc13-C1^HK^ mutants

Cells were stimulated using a paired-pulse protocol consisting of two brief (300 μs) depolarizing pulses at 9 V with a 10 ms interval between them, repeated 10 times at 0.1 Hz. This sequence was performed at progressively increasing extracellular Ca^2+^ concentrations of 0.4, 0.75, 1.5, 3, and 6 mM. The procedure began with a bath volume of 2 ml 0.4 mM Ca^2+^ HL3 and after each measurement, 1 ml of this solution was removed and replaced with 1 ml of 1.1, 2.25, 4.5 or 9 mM Ca^2+^ HL3, respectively, to reach the target concentrations. The solution was gently mixed with a pipette, and the preparation was given a 1-minute acclimation period before the subsequent recording. Paired-pulse ratios (PPRs) were determined by dividing the amplitude of the second evoked excitatory postsynaptic current (eEPSC_2_) by the first (eEPSC_1_) and averaging these values across all 10 stimulations for each cell. Adapted from^66^.

#### Phorbol ester - electrophysiology (PMA, Fig. 6)

For PMA experiments 2 µM PMA (2 mM stock in DMSO, stored at −20°C) or the same volume of DMSO for control experiments was mixed with Ca^2+^-free HL3. 40 µl of the respective solution were incubated on the opened filet for 10 minutes at RT and immediately afterwards three times rinsed with Ca^2+^-free HL3 before transferred to the bath chamber. TEVC recordings were performed as described above in HL3 + 0.4 mM Ca^2+^.

### Single particle tracking PALM

Live sptPALM experiments were performed on male 3^rd^ instar larvae of Unc13-mEOS3.2 and/or Unc13-C1^HK^-mEOS3.2 flies at 25°C (Fig. 2, Fig. S2.2, Fig. S2.3, Fig. 5 and Fig. S5.1). Larval body wall preparation for single particle tracking were prepared according to previously described protocols^11 108^. Imaging experiments were carried out using an inverted total internal reflection fluorescence (TIRF) setup. The microscope (Nikon Eclipse Ti) had a 100x NA 1.49 Apo TIRF oil objective (Nikon). Up to 10.000 images were recorded at a frame rate of 20 Hz using an EMCCD camera (iXon+ 897, Andor Technology) controlled by NIS-Elements (Nikon). Larval NMJ boutons were focused on the live HRP-488 staining used as a reference for the Z-plane. A 1.6 magnification lens was employed to achieve a final pixel size of 107×107 nm.

Male larvae of endogenously tagged Unc13-mEOS3.2 (Fig. 2, Fig. S2.2, Fig. S2.3, Fig. 5 and Fig. S5.1) and/or the Unc13-C1^HK^-mEOS3.2 (Fig. 5 and Fig. S5.1) were prepared, with or without octopamine live drug applications, and were ^11^￼. Male mutant and control larvae were dissected in in Ca^2+^ and Mg^2+^ free HL3.1 saline (in mM: NaCl 70, KCl 5, NaHCO3 10, Sucrose 115, Trehalose 5, HEPES 5). Afterwards, larvae were briefly rinsed in HL3.1 imaging buffer containing 4 mM Mg^2+^ and 1.5 mM Ca^2+^ or for octopamine/PMA treatment experiments 10 mM Mg^2+^ and 0.4 mM Ca^2+^. Larvae were subjected to a 5-minute live HRP-488 stain in the imaging buffer and were then imaged at the HRP Z-plane of type 1b NMJs (from segments A2-A4) on muscle 4. Live sptPALM imaging of Unc13 molecule populations at NMJ muscle 4 Ib boutons was performed and imaged for 8.5 minutes per image at an acquisition rate of 20 Hz.

#### Live octopamine drug application in sptPALM experiments

Male larvae of endogenously tagged Unc13-mEOS3.2 (Fig. 2, Fig. S2.2, Fig. S2.3, Fig. 5 and Fig. S5.1) and/or the Unc13-C1^HK^-mEOS3.2 (Fig. 5 and Fig. S5.1) were dissected as described above and live stained for 5-minute HRP-488 stain in the imaging buffer containing DMSO in HL3.1 imaging buffer with 10 mM Mg^2+^ and 0.4 mM Ca^2+^. These larvae were imaged first for an internal control NMJ at the HRP Z-plane of type 1b NMJs (from segments A2-A4) on Muscle 4. Next, a new Muscle 4 type 1b, NMJ is focused on and prepared for imaging. Before imaging, DMSO containing HL3.1 imaging buffer is replaced with 20 µM octopamine containing HL3.1 imaging buffer and pumped under the filet for 30 seconds and imaged at the prepared and focused NMJ. All imaging was performed live sptPALM imaging of Unc13 molecule populations at NMJ muscle 4 Ib boutons and imaged for 8.5 minutes per image at an acquisition rate of 20Hz.

#### Analysis of stepwise mEOS Photobleaching

A subset of the same raw data analyzed in Figures 2 and S2.3 and Figures 5 and S5.1 data, is used for a stepwise bleaching of the Unc13 attached fluorophore mEOS3.2. Before molecule counting and localization detection, the movies were drift-corrected and cropped to exclude movement artefacts, employing the ImageJ plugins NanoJ ^109^ or Thunderstorm ^110^. Analysis of the local channel density within confined regions was performed by cluster analysis based on Voronoï tessellation constructed from localized channels employing the software package SR-Tessler ^111^. Localization and trajectory reconnection of mEOS signals were performed via a wavelet-based algorithm and a simulated annealing algorithm as previously described, which considers molecule localization and total intensity. To prevent incorrect reconnections between trajectories, all sub-trajectories were analyzed as individual trajectories. Diffusion coefficients (D) were derived from the exponential fitting of the first eight points of the MSD plots. MSD plots of immobilized molecules (on fixed samples) revealed that, under our imaging conditions, molecules with D ≥ 0.002 μm2/s can be considered as mobile. The exponential fit of the MSD determined the radius of confinement. Trajectories with radius of confinement outside the 800 nm-70 nm range were excluded from the analysis. Tracking of individual Unc13 molecules was done under these parameters using PALMtracer plugin of Metamorph^112^.

#### Tessellation Analysis of Unc13 localizations within AZ cluster and Nanocluster distributions from sptPALM images

SR-Tessler software and the ImageJ NanoJ-SRRF were employed to process and segment each Unc13 localization recorded by live sptPALM of all mutant/treated and control images. Tessellation was performed on sptPALM movies which were NanoJ-SRRF drift corrected. The corrected datasets were then subjected to ThunderSTORM analysis to extract all PALM localizations of Unc13 molecules in an image containing an NMJ with several boutons, with each bouton containing high density localizations of Unc13 at AZs. This extracted map of all Unc13 localizations was imported and processed with SR-Tesseler software to generate a tessellated map of all Unc13 localizations. These analyses parameters were repeated as performed in the first seminal paper to study live endogenous Cacophony localizations^11^, which in turn based its analysis criteria off of localization parameters of mono- and polygonal antibody signal clusters in fixed tissue targeting either BRP (QPAINT) ^113^, the GFP tagged Cacophony (dSTORM) ^8^ and the GFP tagged Un13 molecules (dSTORM) ^86^.

A specified set of parameters was employed to select and analyze the highest-density Unc13 localizations at AZs that reflected the optimal TIRF Z focus and exemplified planar Unc13 localization clusters from the tessellation map of all Unc13 channel localizations in an image automatically and robustly. This automated filtration process allowed exclusion of the out of focus and side-view orientations of synapses and their Unc13 clusters.

Tessellation parameters for AZ boundary included: Voronoi object: 100-200 density factor, min Area: 2000, Min Localization: 100-200 NC boundary settings: Voronoi Nanocluster object: 1.5 density factor, min Area: 50, Min Localization: 20. List of AZ clusters and their nanoclusters: size, area, number of localizations, density, and number of clusters were exported and filtered to remove all AZ clusters missing a nanocluster and also clusters outside 600-20 nm AZ and Nanoclusters limits. Furthermore, we applied limitations to Unc13 localization number= AZ boundary: 3000 NC boundary: 3000, diameter size=AZ boundary: 600-20 nm. NC boundary: 300-20 nm, Area: AZ boundary: 50000 nm^2^. NC boundary: 7000 nm^2^ and density=AZ & NC boundary: 1.5 localizations/nm^2^ to define well the nanodomain of Unc13 channels at the presynaptic membrane of the Drosophila NMJ. We took into consideration the parameters set for Unc13 localizations from recent dSTORM and QPAINT imaging studies and our previous study and from recent Unc13 STORM studies ^86^.

#### Quantification and statistical analysis of sptPALM data

The statistical analysis was carried out using Prism software (GraphPad). To test the statistical significance between any two conditions, Kolmogorov-Smirnov test was used to assess the of distribution differences in the diffusion coefficient data, confinement data and tessellation data (Fig. 2, Fig. S2.2, Fig. S2.3, and Fig. S5.1). sptPALM data analysis for mobility, tessellation, well analysis, and channel numbers are shown either as median and an IQR or as mean ± SEM, as described in their respective figure legends. Channel numbers in Fig. S2.3 were tested with paired (Fig. S2.3) and unpaired (Fig. S5.1) T-test U test. Data are presented throughout the text as mean ± SEM or the median with the IQR as indicated. Significance levels are given as *P < 0.05; **P < 0.005; ***P < 0.0005; ****P < 0.0001; and ns, nonsignificant.

### Fluorescence imaging and analysis of AP-induced presynaptic Ca^2+^ influx

For the estimations of Ca^2+^ influx in presynaptic terminals of the *Drosophila* larval NMJ, we used the fluorescent Ca^2+^ sensor GCaMP8f fused to mScarlet and the synaptic vesicle protein Synaptotagmin ^101^, under OK6-Gal4 as a motoneuron-specific driver *(;OK6-Gal4/+;UAS-syt-mScarlet-GCaMP8f/+*). We dissected third instar larvae, exposing the larval body wall with intact motor nerves, in HL-3 saline on a Sylgard-184® (Sigma-Aldrich, St. Louis, MO, USA) block. Dissected larvae were then transferred to the recording chamber containing 2ml of HL-3 with 0.4 mM Ca^2+^ and efferent motoneuron axons enervating the muscle 4 (lateral longitudinal muscle; LL1) were sucked into a polished glass electrode containing a chlorided silver wire, controlled via manual micromanipulators by a patch electrode holder (ALA Scientific Instruments, NY, USA) connected to an isolated stimulator (model ISO-STIM-01D, NPI electronics, Germany). Recordings were performed on the µManager 2.0 software (https://micro-manager.org) through epifluorescence microscopy (Olympus BX51WI, Olympus, Japan) using a white-light LED source with GFP filter (model pE-300^white^, CoolLED, England), an Olympus LUMFL 40X 0.8 NA water-dipping objective, and a Hamamatsu Orca-flash 4.0 LT3 digital camera (Hamamatsu Photonics, Japan) at 5 Hz framerate and 100 ms exposure. Imaging of muscle 4 Ib NMJs from abdominal segments 2 to 4 was conducted over 10 s, and at 5 s, 20 stimuli were applied to the nerve at 20 Hz in 300 µs 7V depolarization pulses. Synchronization of video recording initiation and stimulation timing was executed using the Axon^TM^ Digidata 1550B low-noise data acquisition system and Clampex 11.3 software (Axon Molecular Devices, CA, USA). For naïve controls (Fig. S1), two identical recording sessions were performed 2 minutes apart from each other. For the octopamine pharmacological intervention (Fig. 1D), 20 µM ice-cold DMSO-diluted octopamine (Sigma-Aldrich) was added to the HL3 in the recoding chamber (0.001% DMSO final concentration) 1 minute after the first recording session and incubated for an additional 1 minute before the second recording session. 0.1 M spermine (dilution of 1 µl H_2_O per ml of HL3, Sigma-Aldrich) was added to HL3 to avoid muscle contraction during recordings.

Analysis of GCaMP8f fluorescence was performed using Fiji-ImageJ v1.54f. We manually selected a ROI around the basal fluorescent signal of the NMJs (Fig. S1.1D) and measured the mean pixel gray values within the selection over time. ROIs of identical size, shape, and location were used in the two recording sessions. For each recording, background fluorescence from each frame was measured in a region of the same size and shape outside of the NMJ and subtracted from the signal. Quantification was then performed by subtracting the average fluorescence 1s before the stimulation (F_0_) from the fluorescence signal of the trace for each frame (ΔF), and then normalized by the same baseline fluorescence F_0_, yielding the measurement (ΔF-F_0_)/ F_0_. As abnormal basal levels of synaptic Ca^2+^ influx may indicate poor neuronal health, we identified and excluded outlier fluorescence signals within the first recording sessions by the ROUT Method (Q=1%) using the GraphPad Prism v10.3.1 software. Two NMJ recordings, one from each experiment (with and without octopamine incubation), were excluded using this method.

### Behavior data collection and data analysis

#### Crawling assay

3^rd^ instar larvae were grown at 25°C/65% humidity in 12-hour light/12-hour dark condition, larvae were kept at low density in standard food vials. The experiments were conducted at 22°C. Individual larvae were carefully placed in the center of a 14 cm petri dish filled with 1% agarose. Crawling behavior was recorded for 1 minute at 15 frames/s with the Rapsberry Pi virtual reality setup (PiVR) ^114^. Then, larvae were maintained in food-free agar petri dishes for 2h and subjected again to the crawling assay. To display trajectory plots individual trajectories were centered. The crawling speed was computed with scripts described in ^114^. The statistical analysis was performed in R (R Core Team (2024). https://www.R-project.org/) For statistical comparisons, a two-sample t-test was employed. Additionally, a pairwise comparisons t-test was used to analyze the differences between the fed and starved conditions. *n* represents one animal.

#### Adult locomotor activity assay

Adult locomotor activity was recorded by *Drosophila* Activity Monitors (DAM5H) from TriKinetics. Adult flies raised at controlled densities through larval stages and aged 3-5 days post eclosion were anesthetized with CO_2_ and individually loaded into 5mm x65 mm plastic tubes. The tubes were sealed at one end with a cut foam plug and in the other with a 0.2 mL Eppendorf tube containing food (5% sucrose, 5% yeast, 1% agar) which was exchanged every 2-3 days. The assembled monitors were maintained on a 12-hour light/ 12-hour dark cycle at 25 °C with 65% humidity. The number of moves (when a fly enters a beam after exiting another) from each tube was uploaded every 30 seconds and used as a proxy for locomotor activity. Data extracted within the first 24 hours of monitoring was excluded due to acclimatation of flies to the environment. Data extraction and analysis was performed using DAMSystem3 software (TriKinetics) and a custom python script.

#### Fecundity and egg viability assays

Adult flies raised at controlled densities through larval stages and aged 4-6 days post eclosion were anesthetized with CO_2_ and transferred into embryo collection chambers (15 females and 5 males per chamber). The chambers rested on top of a petri dish containing apple juice agar media (33 % apple juice, 2 % sucrose, 2,6% agar) covered with a thin layer of yeast paste on which the flies deposit eggs. The chambers were left on a 12-hour light/ 12-hour dark cycle at 25 °C with 65% humidity for 24 hours. Fecundity was estimated by manually counting the number of eggs deposited on each dish under a stereo microscope (Olympus SZ51) divided by the number of females remaining in its respective chamber. Egg viability was estimated by transferring groups of 50 eggs to vials with mashed fresh food. The eggs were then allowed to develop and after 10 days the number of viable pupae was counted.

#### Longevity assay

Adult flies raised at controlled densities through larval stages and aged 4-6 days post eclosion were anesthetized with CO_2_ and separated by gender in groups of 20 per replicate. The number of dead flies was noted every 2-3 days. The remaining flies were transferred into fresh vials with new food every 2-3 days throughout the assay.

## Acknowledgements

This work was supported by grants from the Deutsche Forschungsgemeinschaft (DFG) to A.M.W. (Emmy Noether Programme, project ID 261020751 and Transregio SFB 186, project ID 278001972), to M.H. (HE3604/11-1), the Novo Nordisk Foundation (Young Investigator Award project ID NNF19OC0056047 to A.M.W.). We thank Mathias Böhme for help in the planning of the Unc13 C1 and C2B domain mutants. We thank Dion Dickman for sharing the *UAS-syt-mScarlet-GCaMP8f* flies. We also thank Matthieu Louis for the PiVR system. Stocks obtained from the Bloomington Drosophila Stock Center (NIH P40OD018537) were used in this study.

## Author contributions

A.M.W. conceived the project. A.M.W., S.J.S., M.H. and U.T. supervised the project. N.B., Ti.G., To.G., K.S.C., M.B., C.F.C., T.C.M., and L.C. generated experimental data. N.B., Ti.G., To.G., K.S.C., M.B., C.F.C., T.C.M., L.C., H.K. and M.H. analyzed the data. All authors interpreted the data, contributed to the writing and critically reviewed the article.

