## Supplementary_Figures_and_Legends for "Monoamine-induced diacylglycerol signaling rapidly accumulates Unc13 in nanoclusters for fast presynaptic potentiation"

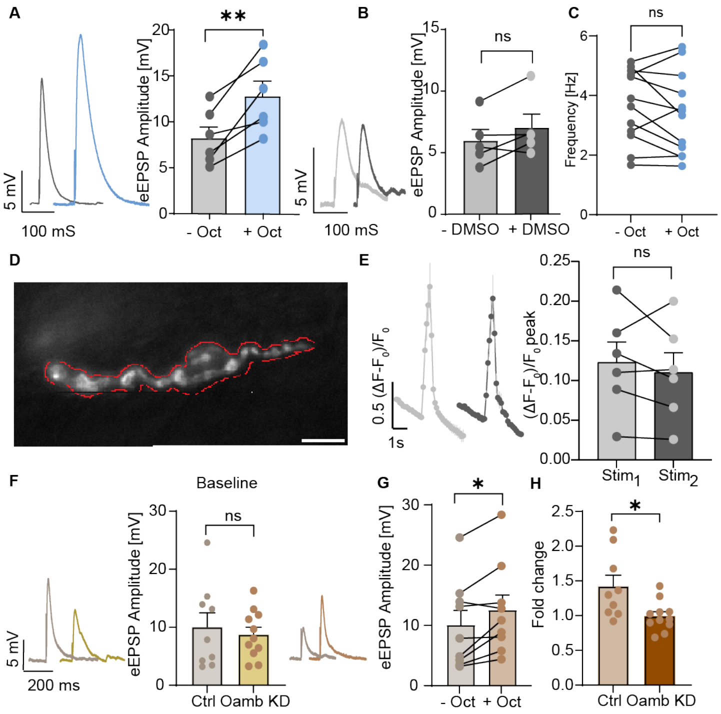


**Figure S1.1: 1-minute incubation time alone has no effect on synaptic transmission.**
(**A, B, C, F, G, H**) Analysis of current clamp recordings (muscle 6 NMJs, (**C, E, F, G, H**) 0.4 mM or (**A** and **B**) 0.3 mM extracellular $Ca^{2+}$) of AP-evoked synaptic activity. (**A**) Representative AP-evoked eEPSP responses (average of five repetitions in one cell) from control synapses (w1118) before (light grey) and after 1-minute 20 µM octopamine incubation (blue) and quantification of eEPSP amplitudes. (**B**) Representative AP-evoked eEPSP responses (average of five repetitions in one cell) from control synapses (w1118) before (grey) and after 1-minute DMSO incubation (dark grey) and quantification of eEPSP amplitudes. (**C**) Analysis of current clamp recordings (muscle 6 NMJs, 0.4 mM extracellular $Ca^{2+}$) of the frequency of spontaneous activity (mEPSP) in wildtype animals before (-oct, grey) and one minute after (+oct, light blue) 20 µM octopamine treatment. (**D**) Example frame of basal fluorescence ($F_0$) of presynaptic GCaMP8f (0.4 mM extracellular $Ca^{2+}$, 100 milliseconds exposure). Red line indicates the ROI used for read-out of the fluorescence signal. (**E**) Quantification of GCaMP8f fluorescence signals from muscle 4 NMJs in naive controls before (dark grey dots, -oct) and after (light grey dots, +oct) a 2-minute interval between recording sessions (represented as averages and SEMs of each recording frame) with paired comparison of the peak florescence signal between sessions. (**F**) Right: representative AP-evoked eEPSP responses (average of five repetitions in one cell) from Ctrl (+>UAS-OAMB-RNAI) (grey) and OAMB KD (*OK6-Gal4>UAS-OAMB-RNAi*) (yellow) mutant synapses and quantification of eEPSP amplitude. (**G**) Representative AP-evoked eEPSP responses (average of five repetitions in one cell) from control synapses before (grey) and after 1-minute 20 µM octopamine incubation (beige) with quantification of eEPSP amplitudes. (**H**) Comparison of the fold-change in eEPSP amplitudes after 1 minute of octopamine incubation divided by the eEPSP amplitudes prior to treatment for the control condition (beige) and OAMB KD animals (brown). Number of cells (n) and animals (N) investigated: n/N(Octopamine, A) = 6/6; n/N(DMSO, **B**) = 5/5; n/N (w1118 frequency, **C**) = 12/12; n/N(Syt::GCaMP8f, **E**) = 6/6; n/N(OAMB ctrl, **F,H**) = 9/9;

n/N(OAMB KD, **G,H**) = 11/11. For exact genotypes see methods. Data depict mean values ± SEM. Statistical analysis with paired parametric t-tests (**A, B, C, E, G**) or Mann-Whitney test (**F, H**). n.s., p > 0.05; *p % 0.05; **p % 0.01. Scale bar: 10 µm

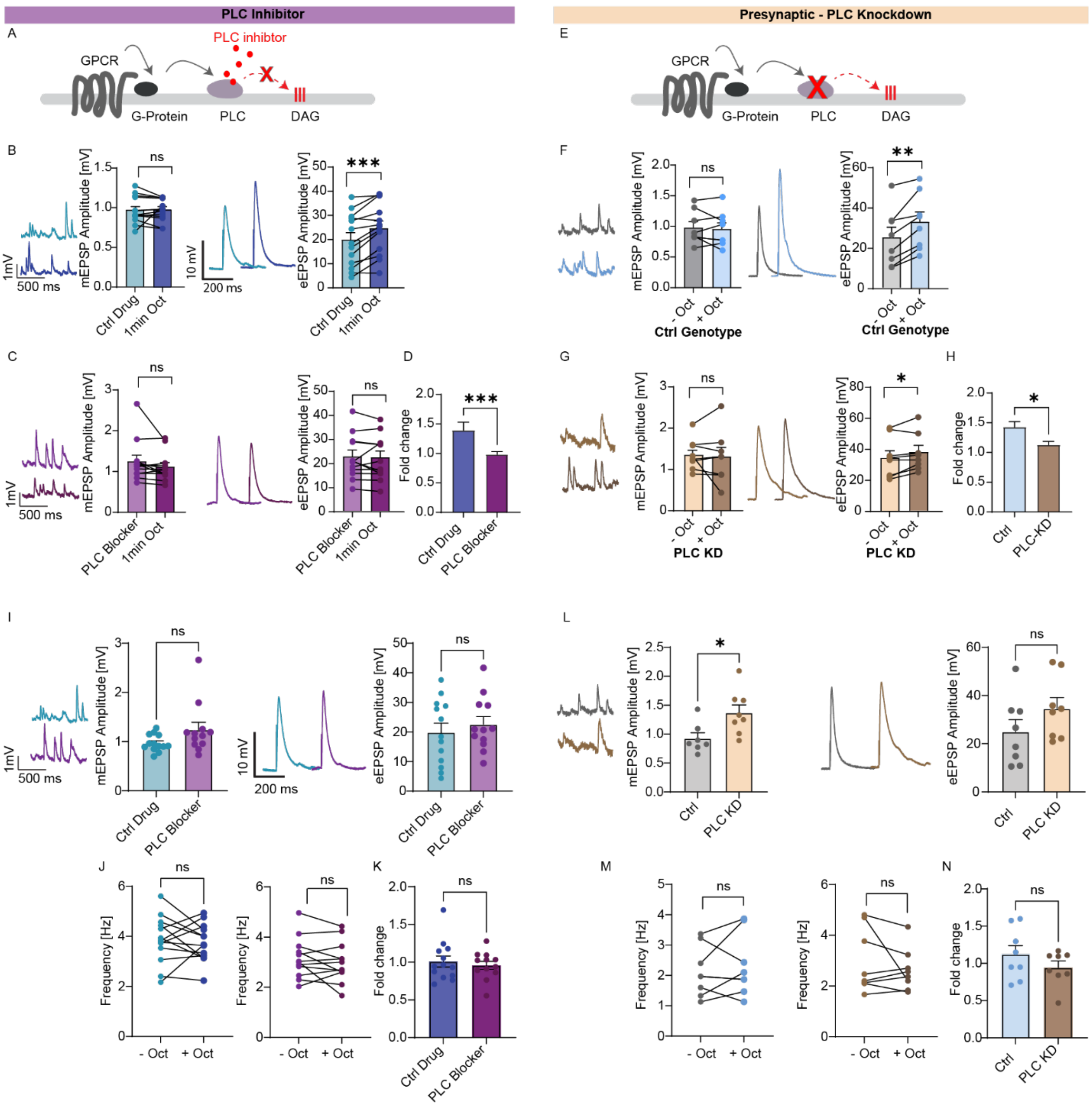


**Figure S1.2: Octopamine-induced potentiation requires presynaptic phospholipase C.** (**A** and **E**) Scheme of phospholipase C signaling pathway indicating a G protein-coupled receptor (GPCR) g-protein, Phospholipase C (PLC) and diacylglycerol (DAG) and the PLC inhibitor blocking PLC (**A**) or the presynaptic PLC knockdown (**E**). (**B**, **C**, **D**, **F**, **G**, **H**, **I**, **J**, **K**, **L**, **M**, **N**) Analysis of current clamp recordings (muscle 6 NMJs, 0.4 mM extracellular $Ca^{2+}$) of spontaneous (left) and AP-evoked synaptic activity (right). (**B**) Left: representative example traces of spontaneous mEPSPs from Ctrl animals (w1118) with PLC control drug (U73343, 1 µM; light blue) and after 1 min 20 µM octopamine (dark blue) and quantification of mEPSP amplitudes after PLC ctrl drug before octopamine (light blue: ctrl drug, -oct) and after 1 minute incubation with 20 µM octopamine (+oct, dark blue) Right: representative AP-evoked eEPSP responses (average of five repetitions in one cell) Ctrl animals (w1118) with 1-minute PLC ctrl drug incubation (U73343, 1 µM; light blue) and followed by 1-minute incubation with 20 µM octopamine (dark blue) and quantification of eEPSP amplitudes. (**C**) Left: representative example traces of spontaneous mEPSPs from Ctrl animals (w1118) with PLC inhibitor (U73122, 1 µM; light purple) and followed by 1-minute incubation with 20 µM octopamine (dark purple) and quantification of mEPSP amplitudes with PLC inhibitor and before octopamine (light purple) and after 1-minute incubation with 20 µM octopamine (dark purple). Right: representative AP-evoked eEPSP responses (average of five repetitions in one cell) in Ctrl animals (w1118) with 1 minute PLC inhibitor (U73122, 1 µM; light purple) and followed by 1-minute incubation with 20 µM octopamine (dark purple) and quantification of eEPSP amplitudes (Panel C (right) reused from Fig. 1C). (**D**) Comparison of the fold change in eEPSP amplitudes after and before 1 minute of octopamine incubation between control drug (blue) and PLC inhibitor (purple). (**F**) Left: representative example traces of spontaneous mEPSPs from control genotype synapses before and after 1-minute octopamine incubation (-oct, grey; +oct, blue) and quantification of mEPSP amplitudes. Right: representative AP-evoked eEPSP responses (average of five repetitions in one cell) from control genotype synapses and quantification of eEPSP amplitude (-oct, grey; +oct, blue). (**G**) Left: representative example traces of spontaneous mEPSPs from PLC knockdown synapses before octopamine incubation (beige, top) and after 1-minute incubation with 20 µM octopamine (brown, bottom) together with a cell-wise quantification of mEPSP amplitudes before (-oct, beige) and 1 min after incubation with 20 µM octopamine (+oct, brown). Right: representative AP-evoked eEPSP responses (average of five repetitions in one cell) from PLC knockdown synapses and quantification of eEPSP amplitude (-oct, beige; +oct, brown). (**H**) Comparison of the fold change in eEPSP amplitudes after 1 minute of octopamine incubation between Ctrl genotype (blue) and PLC knockdown (brown) animals. (**I**) representative example traces of spontaneous mEPSPs after 1- minute control drug incubation (1 µM, blue, top) and after 1-minute PLC inhibitor treatment (1 µM, purple, bottom) together with a cell-wise quantification of mEPSP amplitudes after 1-minute control drug (blue) and PLC inhibitor (purple) incubation. Right: representative AP-evoked eEPSP responses (average of five repetitions in one cell) from Ctrl synapses with displayed drug treatment and quantification of eEPSP amplitudes after indicated drug treatment (ctrl drug=blue; PLC inhibitor=purple). (**J**) Quantification of mEPSP frequency in control condition (left) PLC inhibitor condition before and after one-minute 20 µM octopamine incubation (right) with the comparison of the fold change in mEPSP frequency between control drug and PLC blocker (**K**). (**L**) Left: representative example traces of spontaneous mEPSPs from Ctrl (grey, top) and PLC knockdown (beige, bottom) synapses and quantification of mEPSP amplitudes. Right: representative AP-evoked eEPSP responses (average of five repetitions in one cell) from Ctrl and PLC knockdown synapses and quantification of eEPSP amplitudes. (**M**) Quantification of mEPSP frequency in control genotype (left) and PLC KD before and after one-minute 20 µM octopamine incubation (right) with the comparison of the fold change in mEPSP frequency between control genotype and PLC KD (**N**). Number of cells (n) and animals (N) investigated: n/N: n/N(*Ctrl drug*) = 13/13, n/N(PLC inhibitor) = 12/12, n/N(*Ctrl genotype*) = 8/8, n/N(PLC knockdown) = 8/8. For exact genotypes see methods. Data depict mean values ± SEM. Statistical analysis with unpaired, Mann-Whitney U test (**D, H, I, L, K, N**), or with paired parametric t-tests (**B, C, F, G, J, M**). n.s., p > 0.05; *p % 0.05; **p % 0.01; ***p % 0.001.

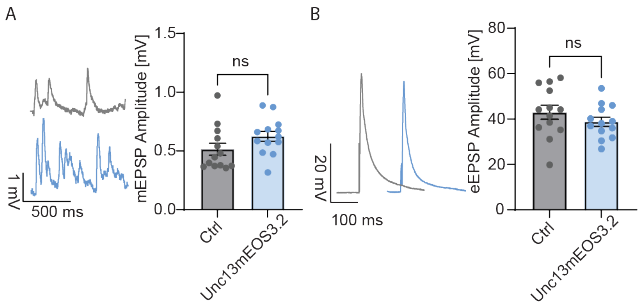


**Figure S2.1: CRISPR Cas9 mediated mEOS3.2 tagging of the endogenous Unc13 protein does not affect baseline synaptic properties.** (**A**) Representative example traces of spontaneous mEPSPs from control (Ctrl) animals (wildtype, *w1118*; grey, top) and Unc13mEOS3.2 animals (blue, bottom) together with a cell-wise quantification of mEPSP amplitudes. (**B**) Representative AP-evoked eEPSP responses (average of five repetitions in one cell) from control synapses (grey) and Unc13mEOS3.2 animals (blue) and quantification of eEPSP amplitudes. Number of cells (n) and animals (N) investigated: n/N(*Ctrl*) = 13/13; n/N(Unc13mEOS3.2) = 13/13. For exact genotypes see methods. Data depict mean values ± SEM. Statistical analysis with unpaired t-tests. n.s., p > 0.05.

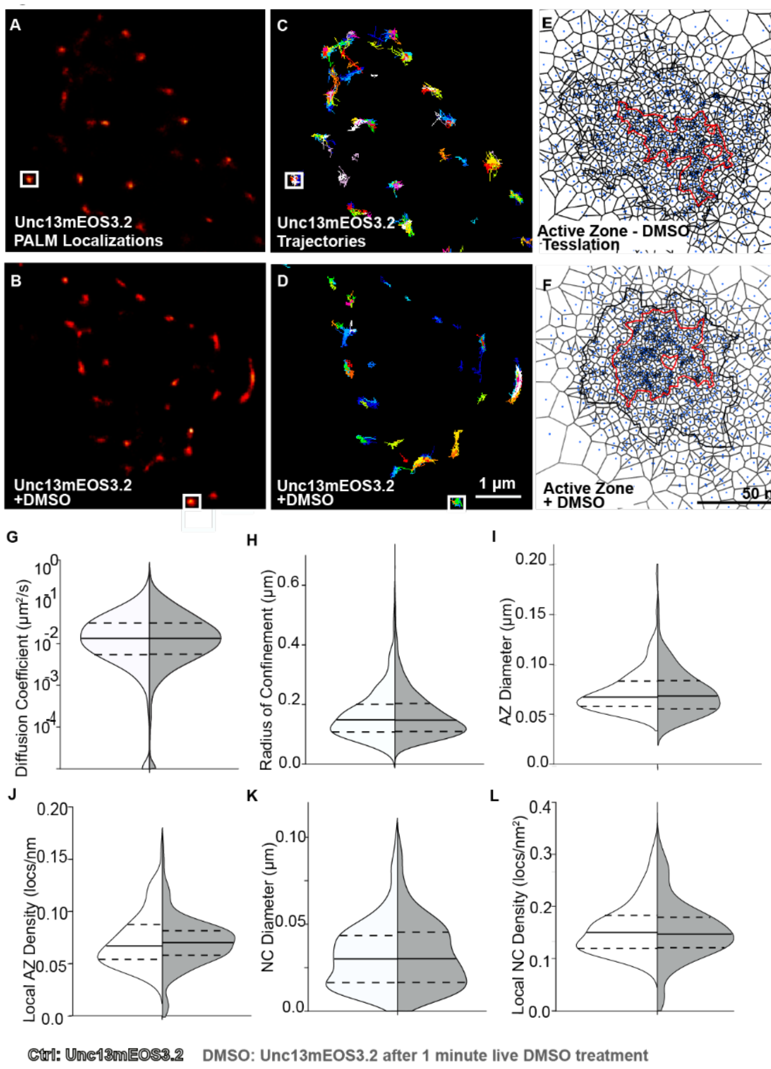


**Figure S2.2: Live 1-minute application of DMSO does not impact *in vivo* imaging of endogenously tagged Unc13 molecules.** Negative controls in this figure were done concomitantly with experimental conditions in Figure 6. Live sptPALM imaging of Unc13mEOS3.2 at muscle 4 NMJs, performed in 0.4 mM $Ca^{2+}$ and 10 mM $Mg^{2+}$ containing HL3.1, before (Internal Control/Ctrl) and after a 1-min incubation of DMSO in HL3.1 (DMSO). Images show representative sptPALM recordings (**A** and **B**), trajectory maps (**C** and **D**), and tessellation analysis representations of Unc13mEOS3.2 before and after 1-minute DMSO treatment (**E**, and **F**). (**G** and **H**) Quantification of diffusion coefficients and radii of confinement from live Unc13 channel sptPALM imaging. Ctrl NMJs: 1437 single-molecule trajectories; 1-minute DMSO treatment = 2966 single-molecule trajectories. (**I** to **L**) Tessellation analysis from the same sptPALM dataset was analyzed in (**F**) to (**I**) for diameters and densities of Unc13 localizations within AZ (**I**—**J**) and NC cluster (**K**—**L**) boundaries. Here, 10 animals per condition generated 252 AZs (control) and 201 AZs (DMSO), respectively, that were analyzed. Number of NMJs (n), animals (N), number of AZs (X) and number of individual trajectories (**Y**) investigated: n/N/X/Y(*Ctrl*, A) = 10/10//252/1437; n/N (DMSO, B) = 10/10/201/2966; Data depict mean values ± SEM. Statistical significance is denoted as asterisks: **P < 0.01, ***P < 0.001, and ****P < 0.0001. Data distribution was statistically tested with a Kolmogorov-Smirnov test. AU, arbitrary units. Scale bars, 1 µm (**A** to **D**) and 50 nm (E and F).

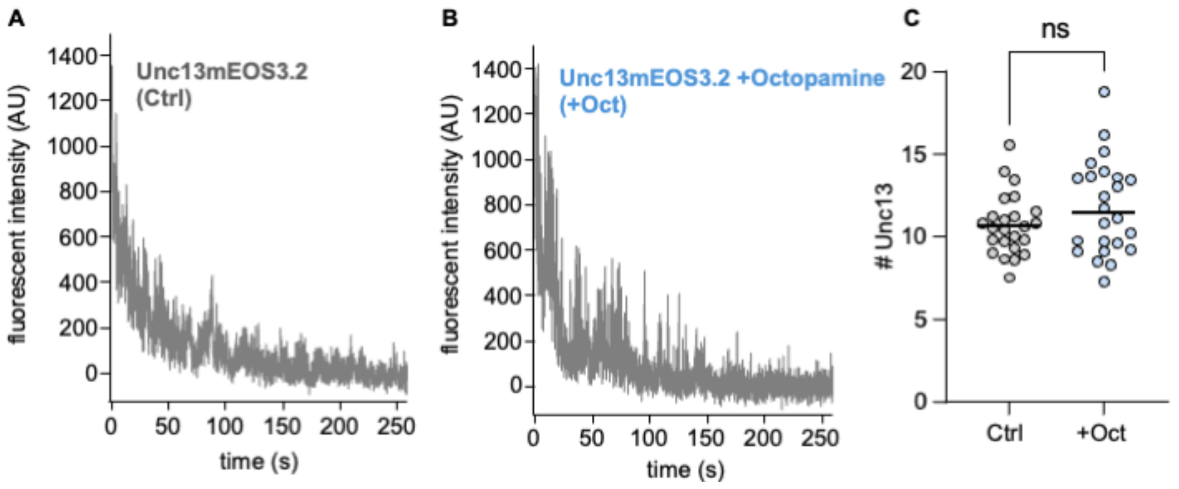


**Figure S2.3: Molecular counting of Unc13mEOS3.2 sptPALM live imaging experiments in *Drosophila* larvae.** Estimation of relative channel numbers based on the blinking fluorescent emission of mEOS3.2 tagged UNC13 molecules via bleach curve analysis. Averaged fluorescent intensity of individual synapses was recorded for 50s and 250s. Fluorescent intensity measurements over time normalized first to the average background signal and second to the average amplitude of single fluorophore events at the end of the recording. Bleach curve fluorescent intensity measurements of (**A**) Unc13mEOS3.2 controls (Ctrl, -Oct), and (**B**) Unc13mEOS3.2 with octopamine (+ Oct). (**C**) Quantification of relative Unc13 numbers from control and after octopamine. For comparison, quantification of relative Unc13 numbers extracted from a subset of the same data analyzed in live sptPALM experiments of Fig. 2. Number of NMJs (n) and animals (N) and number of AZs (X) investigated: n/N/X(*Ctrl in **C***) = *3*/3/24; n/N/X(+Oct *in **C***) = *3*/3/24. Data distribution was statistically tested with paired T-test. Statistical significance is denoted as asterisks: **P* < 0.01,****P* < 0.001, and *****P* < 0.0001

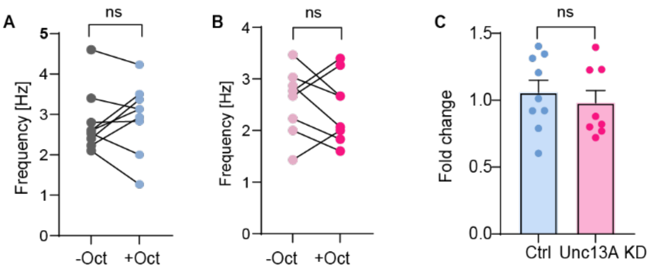


**Figure S3.1: Octopamine incubation does not change mEPSPs frequency in Unc13A KD animals.** Analysis of current clamp recordings (muscle 6 NMJs, 0.4 mM extracellular $Ca^{2+}$) of mEPSP frequency activity. (**A**) Analysis of mEPSP frequency in control animals before (-oct, grey) and one minute after 20 µM octopamine treatment (+oct, light blue). (**B**) mEPSP frequency before (-oct, light pink) and one minute after (+oct, pink) octopamine treatment in Unc13A KD animals. (**C**) Comparison of the fold-change in mEPSP frequency after 1 minute of octopamine incubation divided by the mEPSP frequence prior to treatment for the control condition (blue) and Unc13A KD animals (pink). Number of cells (n) and animals (N) investigated: n/N: n/N(Genetic ctrl) = 9/9; n/N(Unc13A KD) = 8/8. For exact genotypes see methods. Data depict mean values ± SEM. Statistical analysis with unpaired, Mann-Whitney U test (**C**), or with paired parametric t-tests (A and B). n.s., $p > 0.05$.

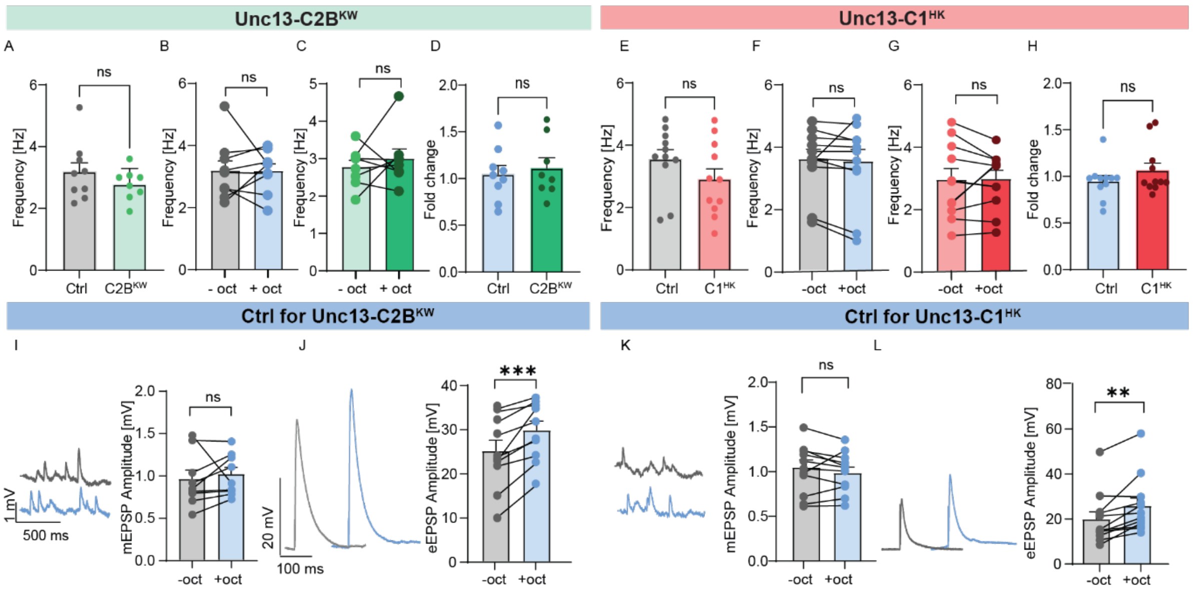


**Figure S4.1: Analysis of mEPSP frequency and control experiment for the gain of function mutants of Unc13's C1 and C2B domain.** (**A**) Comparison of current clamp recordings (muscle 6 NMJs, 0.4 mM extracellular Ca$^{2+}$) shows the frequency of spontaneous activity (mEPSP) in wildtype animals and Unc13-C2B$^{KW}$ mutants before octopamine treatment. (**B**) Analysis of the frequency of mEPSP in wildtype animals before and after one-minute 20 µM octopamine treatment. (**C**) Analysis of the frequency of mEPSP in Unc13-C2B$^{KW}$ mutants before and after octopamine treatment. (**D**) Comparison of the fold change in mEPSP frequency after 1 minute of octopamine incubation between control condition (blue) and Unc13-C2B$^{KW}$ mutants (green).

(**E**, **F**, **G**, **H**) Same analysis as in (**A**, **B**, **C**, **D**) for Unc13-C1$^{HK}$ mutant flies. (**I** and **J**) Shown in Fig.1A before. (**I**) Representative example traces of spontaneous mEPSPs from control synapses (Unc13-C2B$^{KW}$ experiment) before (grey, top) and after 1-minute octopamine incubation (blue, bottom) and quantification of mEPSP amplitudes. (**J**) Representative AP-evoked eEPSP responses (average of five repetitions in one cell) from control synapses (Unc13-C2B$^{KW}$ experiment) and quantification of eEPSP amplitudes before (grey) and after octopamine incubation (blue).

(**K, L**) Same analysis as in (**I**, **J**) for Unc13-C1$^{HK}$ mutant flies. Number of cells (n) and animals (N) investigated: n/N: n/N(*Ctrl*, **A**) = 9/9, n/N(Unc13-C2B$^{KW}$, **B**) += 8/8, n/N(Ctrl, **D**) = 11/11, n/N(Unc13-C1$^{HK}$, **E**) = 11/11, n/N(Ctrl, G) = 9/9, n/N(Ctrl, **H**) = 10/10, n/N(Ctrl, **I**) = 11/11, n/N (Ctrl, **J**) = 12/12. For exact genotypes see methods. Data depict mean values ± SEM. Difference between means was tested with paired parametric t-test (**B, C, I, J, K, L, F, G**) or Statistical analysis with unpaired, Mann-Whitney U test (**A, D, E, H**). n.s., p > 0.05; **p % 0.01; ***p % 0.001.

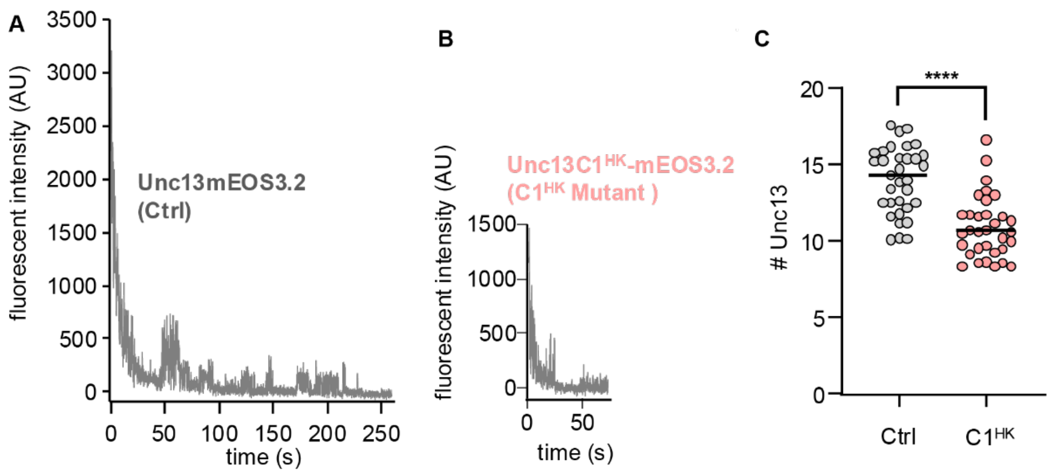


**Figure S5.1: Molecular counting of Unc13-C1$^{HK}$-mEOS3.2 mutant sptPALM live imaging experiments in *Drosophila* larvae.** Estimation of relative channel numbers based on the blinking fluorescent emission of mEOS3.2 tagged UNC13 molecules via bleach curve analysis. Averaged fluorescent intensity of individual synapses was recorded for 50s and 250s. Fluorescent intensity measurements over time normalized first to the average background signal and second to the average amplitude of single fluorophore events at the end of the recording. Bleach curve fluorescent intensity measurements of Unc13mEOS3.2 controls (Ctrl) (**A**), and (**B**) Unc13-C1$^{HK}$-mEOS3.2 mutant (C1$^{HK}$). For comparison (**C**), quantification of relative Unc13 numbers extracted from a subset of the same data analyzed in of the live sptPALM experiments of Fig. 5B-K. Number of NMJs (n) and animals (N) and number of AZs (X) investigated: n/N/X(*Ctrl in C*) = *3*/2/33; n/N/X(C1$^{HK}$) = *5*/2/33. Data distribution was statistically tested with an unpaired T-test. Statistical significance is denoted as asterisks: **P* < 0.01,****P* < 0.001, and *****P* < 0.0001.

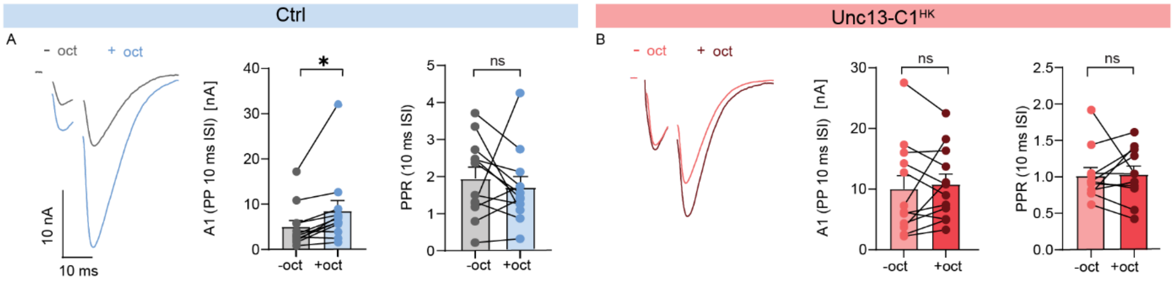


**Figure S6.1: Octopamine acutely potentiates neurotransmitter release in control synapses, but this potentiation is blocked in the Unc13-C1$^{HK}$ mutant.** (**A**) Analysis of control (Ctrl) animals at 0.4 mM extracellular $Ca^{2+}$ concentration. Left: example traces of AP-evoked paired-pulse responses preincubated before 1-minute octopamine incubation (−oct, grey) or after 1-minute 20 µM octopamine incubation (+oct, blue). Middle: quantification of $eEPSC_1$ amplitudes before and after octopamine. Right: quantification of PPR ratios (10 ms interstimulus interval [ISI]). (**B**) Analysis of Unc13-C1$^{HK}$ animals. Left: example traces of AP-evoked paired-pulse responses preincubated before 1-minute octopamine incubation (−oct,light red) or after 1-minute 20 µM octopamine incubation (+oct, dark red). Middle: quantification of $eEPSC_1$ amplitudes before and after octopamine. Right: quantification of PPR ratios (10 ms interstimulus interval [ISI]). Number of cells (n) and animals (N) investigated: n/N: n/N(*Ctrl*, **A**) = 12/12, n/N(Unc13-C1$^{HK}$, **B**) = 12/12. For exact genotypes see methods. Data depict mean values ± SEM. Statistical analysis with paired parametric t-test. n.s., p > 0.05; *p % 0.05; **p % 0.01; ***p % 0.001.

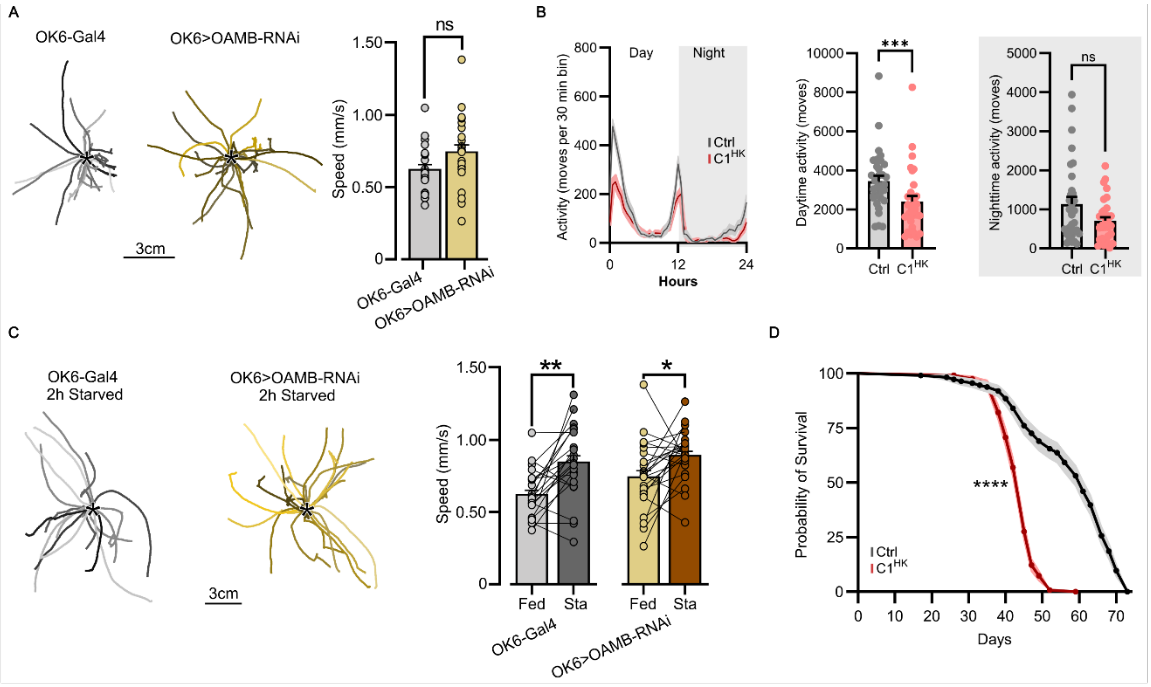


**Figure S7.1: Larvae with OAMB depletion in motor neurons maintain baseline and starvation-induced adaptive crawling speed. Unc13-C1$^{HK}$ males also show decreased locomotion and lifespan. (A)** Trajectories of 1-minute of 3$^{rd}$ instar larvae crawling filmed at 15 frames/s (*, starting position) from OK6-Gal4 control flies (grey traces) and from *OK6-Gal4>UAS-OAMB-RNAi* (light brown traces). Crawling speed is not affected when knocking down the presynaptic OAMB receptors in the motor neurons (OK6-Gal4: n=21; *OK6-Gal4>UAS-OAMB-RNAi*: n=23; two-sample t-test, p=0.052). (**B**) Locomotor activity profile of adult male control (Ctrl) and Unc13-C1$^{HK}$ (C1$^{HK}$) animals across 24 hours (left) and quantification of total activity split in day (centre) and night (right). Number of animals (n): n(Ctrl)=32, n(C1$^{HK}$)=32. Unpaired Mann-Whitney test, daytime activity: p=0.0009, nighttime activity: p=0.1248. (**C**) 2h-starvation significantly increases crawling speed in both groups (*OK6-Gal4>UAS-OAMB-RNAi*: Paired t-test, p=0.016; OK6-Gal4: Paired t-test, p=0.0012). (**D**) Longevity analysis of male control and Unc13-C1$^{HK}$ animals. Lifespan is reduced in male Unc13-C1$^{HK}$ flies. Number of animals (n): n(Ctrl)=113, n(C1$^{HK}$)=123. Log-rank (Mantel-Cox) test, p<0.0001. Error bars represent SEM. * P < 0.05; **P < 0.01; ***P < 0.001.
